# Synergising single-cell resolution and 4sU labelling boosts inference of transcriptional bursting

**DOI:** 10.1101/2022.09.08.506961

**Authors:** David M Edwards, Philip Davies, Daniel Hebenstreit

**Affiliations:** School of Life Sciences, University of Warwick

## Abstract

Despite the recent rise of RNA-seq datasets combining single-cell (sc) resolution with 4-thiouridine (4sU) labelling, analytical methods exploiting their power to dissect transcriptional bursting are lacking. Here, we present a mathematical model and Bayesian inference implementation to facilitate genome-wide joint parameter estimation and confidence quantification. We demonstrate that, unlike conventional scRNA-seq, 4sU scRNA-seq resolves temporal parameters and furthermore boosts inference of dimensionless parameters via a synergy between single-cell resolution and 4sU labelling. We applied our method to published 4sU scRNA-seq data and linked with ChIP-seq data, uncovering previously obscured associations between different parameters and histone modifications.

## Background

The canonical understanding of transcription is that it consists of the steps of initiation, elongation and termination. During initiation of transcription in eukaryotes, RNA polymerase (RNAP) is recruited to the promoter via transcription factors (TF), followed by the synthesis of the first few bases of the new transcript [1]. Elongation succeeds initiation, in which RNAP processes along the gene, incorporating RNA nucleotides into the nascent transcript as it progresses [2]. Upon reaching the transcription end site (TES) termination occurs, in which the transcript and RNAP are released from the DNA [3]. Various processing steps take place at different points during transcription to allow for a mature transcript to be produced, including 5’ capping during initiation, splicing to remove intronic (non-coding) sequences during elongation of protein-coding genes, and polyadenylation and cleavage during termination [1, 2, 3, 4].

Beyond the general mechanism outlined above, transcription is also a stochastic process subject to intrinsic noise through its fundamental dependence on probabilistic collisions between molecules [5, 6], which are often present in relatively low numbers. Additionally, in many cases, transcription occurs only in short, intense bursts of activity followed by prolonged periods of inactivity, resulting in increased cell-cell variability in transcript counts [7, 8]. Indeed, studies have identified a broad spectrum of genes, from those that are transcribed in a Poissonian fashion, such as housekeeping genes, to those which are very bursty in nature and expressed only in relatively short windows [9, 10]. The transcriptional noise and cell-cell variance induced by bursting can be utilised to, for example, achieve alternative cell fates during differentiation of cell populations without requiring explicit control by genetic programming or external signals [11]. There are several different possible mechanisms thought to contribute to bursting, including the process of reinitiation, in which after transcribing a gene the RNAP is immediately recycled to the transcription start site (TSS) instead of simply terminating and disengaging [12]. This requires looping of the gene to bring the TSS and TES into physical proximity [13], and the link between TSS-TES interactions and bursting has been explored recently [14]. The chromatin state of a gene also plays an important role in governing transcriptional bursting dynamics, which in eukaryotes is dictated largely by histone modifications (HM). Different HMs may result in looser or tighter packing of the chromatin, respectively, with the chromatin density around the TSS being correlated with transcriptional noise [15]. Having active HMs at the TSS results in an increased probability of open chromatin, which facilitates initiation. This is proposed to reduce burstiness, possibly by reducing the duration between active periods [15, 16]. More recent studies have also reported genome-wide direct correlations between the presence of specific HMs at gene promoters and general transcriptional noise [17, 18], while further studies have even linked HMs with the underlying bursting dynamics, both at the individual gene level [19] and genome-wide [20]. Transcriptional bursting in bacteria can also result from supercoiling of the DNA [21]. The proposed mechanism is the accumulation of positive supercoiling caused by the RNAP proceeding through the gene, until it reduces the rate of elongation to the point that it prevents further transcription. Intermittent clearing of supercoiling followed by rapid transcription, and subsequent re-accumulation of supercoiling, results in bursty transcription. Studies have also observed the co-condensation of TFs with transcriptional coactivators such as p300, which mediates cooperative activation of genes by clusters of TFs [22]. This cooperative activation results in non-linear gene regulation and increased burst frequency and burst size for genes enriched in coactivators.

Transcriptional bursting may be understood in terms of several parameters (figure 1), including the burst size (transcripts produced per burst, b), burst frequency (bursts per unit time, *κ*), decay rate (transcripts degraded per unit time, *δ*), burst rate (bursts per transcript lifetime, *a* = *κ/δ*) and expression level (mean transcripts per cell, *μ* = *b* × *a*). Many studies make use of fluorescence microscopy-based approaches to interrogate transcriptional bursting dynamics. Single molecule fluorescence in situ hybridisation (sm-FISH) is a particularly popular approach here although the standard procedure offers only a snapshot of transcript counts across a cell population, with no time-variant information. Therefore, the timescales of bursting events may not be discerned [23], allowing estimation of *μ, b* and *a* but not *κ* or *δ*. Some smFISH-based experimental set-ups have progressed towards a level of understanding bursting timescales by using hybridisation specific to nascent transcripts [24, 25], although smFISH approaches generally suffer from scalability. While progress is being made towards multiplexing, it can still only analyse a handful of genes at a time compared with sequencing [26, 27, 28] or requires complex and labourious set-ups [29]. Sophisticated analysis methods [30] have been developed for time-lapse single-cell RNA imaging data [31] which allows dissection of transcription dynamics in great detail, however such approaches are even more limited scale-wise.

**Figure 1:**
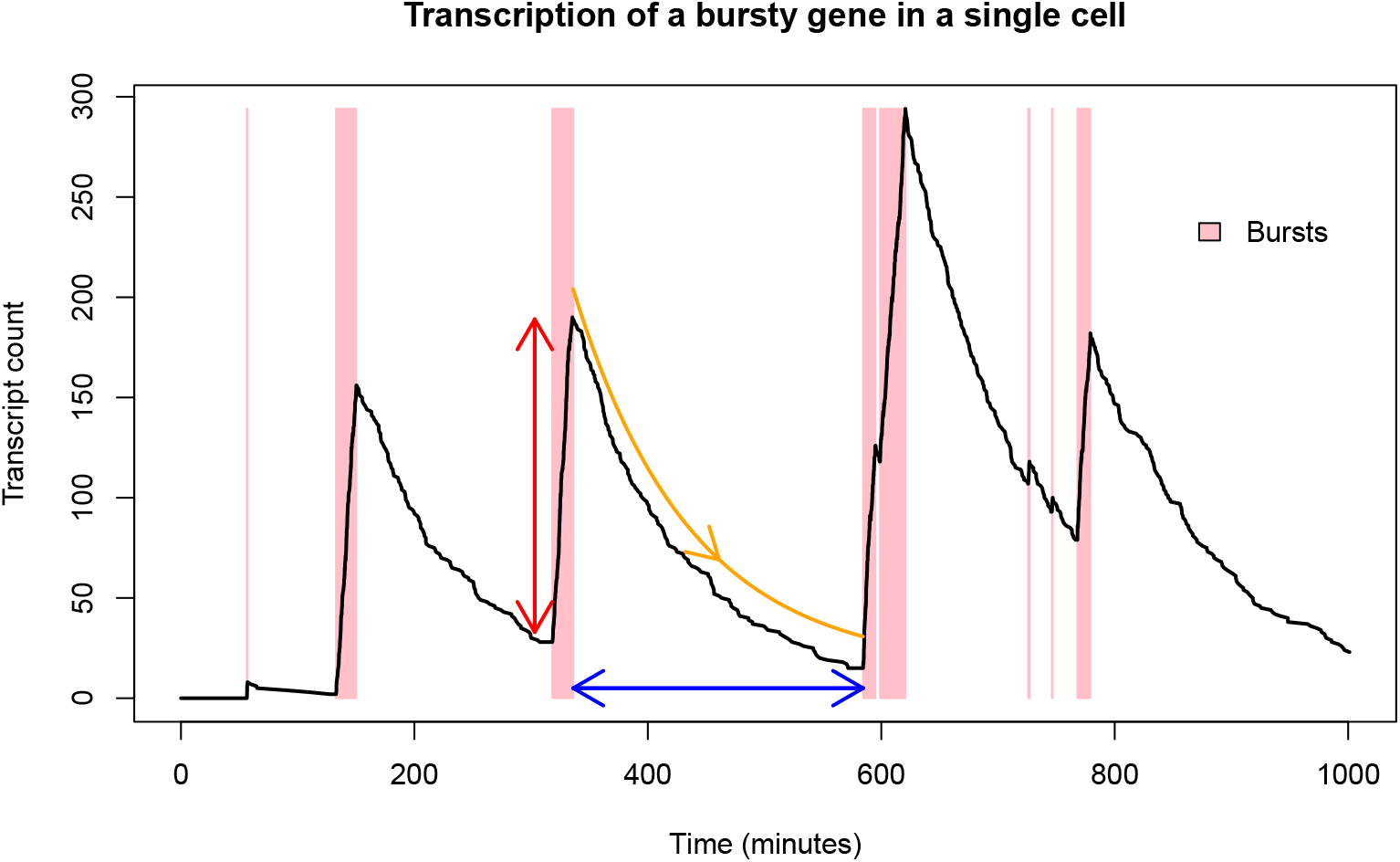
Simulation demonstrating transcriptional bursting for a single gene in a single cell, indicating burst size (red), burst interval (blue, reciprocal of burst frequency), and decay rate (orange, reciprocal of transcript lifetime), while the thickness of the pink shaded regions indicate burst durations.

Single cell RNA-seq (scRNA-seq) experiments are widely used to analyse genome-wide bursting dynamics. However, scRNA-seq suffers from the same issue as standard smFISH regarding analysis of bursting timescales because it only provides a snapshot of the transcriptomes of a population of cells at a single point in time. Therefore, it has only been possible to obtain burst sizes (*b*) and burst rates (*a*), while burst frequencies (*κ*) may not be understood without making assumptions or using prior information on decay rates (*δ*) measured through separate experiments [10, 32, 33, 34]. On the other hand, bulk RNA-seq-based approaches have for several years made use of chemically labelled nucleotides, primarily 4-thiouridine (4sU), to understand RNA synthesis (*b* × *κ*) and degradation (*δ*) rates [35, 36]. The cells are incubated in the presence of 4sU for a given duration, prior to RNA extraction. During this step, 4sU diffuses into the cell nucleus and becomes incorporated into nascently transcribed RNA. Labelled RNA can be bioinformatically distinguished from non-labelled RNA, previously residing in the cell, due to the higher rate of chemically-induced cytosine conversion of 4sU relative to regular uracil. Using mathematical modelling, the ratio of labelled to unlabelled transcripts can be used to estimate the turnover rate [37]. However, since bulk RNA-seq neglects the cell-cell variability, it can not be used to study bursting dynamics. Recent advances combine scRNA-seq with 4sU and such datasets have the potential to fully characterise transcriptional bursting dynamics and their timescales (figure 2). Thus far, they have been used for understanding dynamic changes in the transcriptome and/or RNA turnover/splicing rates that occur throughout the cell cycle and cell state transitions [38, 39, 40, 41, 42]. Studies with data of this type that have looked at bursting have only done so in a limited manner, using empirically-derived statistics as a proxy for burstiness [43], while bursting timescales have remained uncharacterised in recent works [44].

**Figure 2:**
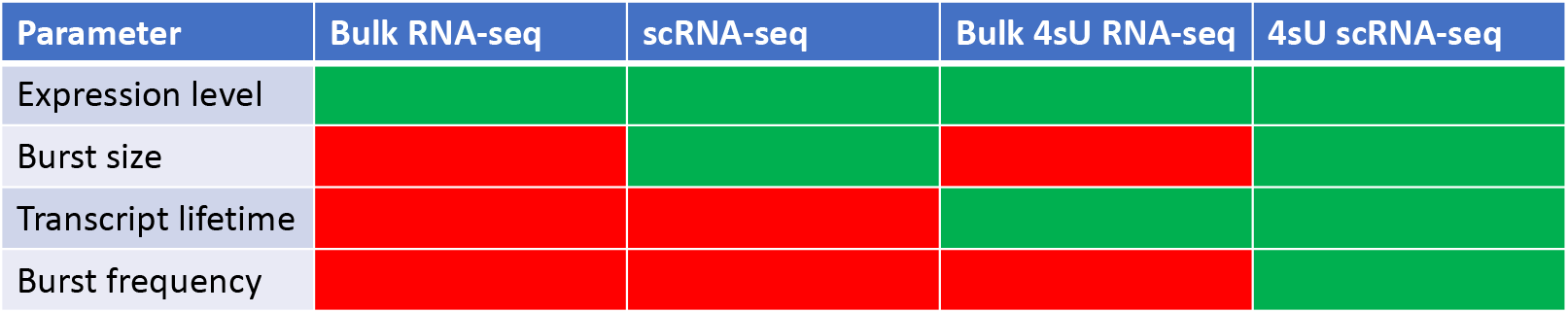
Table showing the parameters governing transcription dynamics that can theoretically be obtained using different RNA-seq approaches with no prior information. Green and red show if a data type does or does not inform a parameter, respectively.

Here we construct mathematical models to relate observables from 4sU scRNA-seq data to the underlying bursting dynamics and develop an adaptive Markov chain Monte Carlo (MCMC) approach for Bayesian inference of the parameters governing those dynamics. We apply this method to published data from [38] and demonstrate that we are able to characterise time-resolved transcriptional bursting dynamics for hundreds of genes in parallel. Our approach generates joint probability distributions of the parameters of interest from which estimates can be extracted and confidence in these quantified. This is the first method for joint inference of time-resolved bursting dynamics on a genome-wide scale and is generally applicable to 4sU scRNA-seq datasets. We also show that, even for the dimensionless parameters which can be obtained with conventional scRNA-seq, the accuracy and reliability of estimates can be improved by incorporating the additional information provided by 4sU scRNA-seq. Finally, we build on a previous study which interrogated correlations between bursting parameter estimates and HMs in a genome-wide manner, linking scRNA-seq with ChIP-seq data [20]. Our analysis reveals position-dependent associations between different bursting parameters and HMs only apparent with 4sU scRNA-seq.

## Results

### Model comparison

We tested the advantages provided by 4sU scRNA-seq data coupled with our inference approach over conventional scRNA-seq by comparing our recovery of known bursting parameter values from a simulated dataset using different likelihood functions (Methods). The MCMC algorithm was run three times, using equations 4, 14 and 15 as the likelihood functions, referred to as L1, L2 and L1+L2.

- L1: The likelihood function of model 1, equivalent to scRNA-seq data without 4sU, relying solely on the UMI counts.
- L2: Equivalent to relying only on single cell T>C conversions, without fully incorporating the UMI counts.
- L1+L2: The likelihood function of model 2, equivalent to 4sU scRNA-seq data, incorporating all of the available information together.

Convergence to the target distribution is shown in figure 3 for each likelihood function, confirming that scRNA-seq data cannot resolve *κ* or *δ*, but does converge for the other parameters, while L2 and L1+L2 converge for all parameters, confirming that 4sU scRNA-seq data can time-resolve bursting.

**Figure 3:**
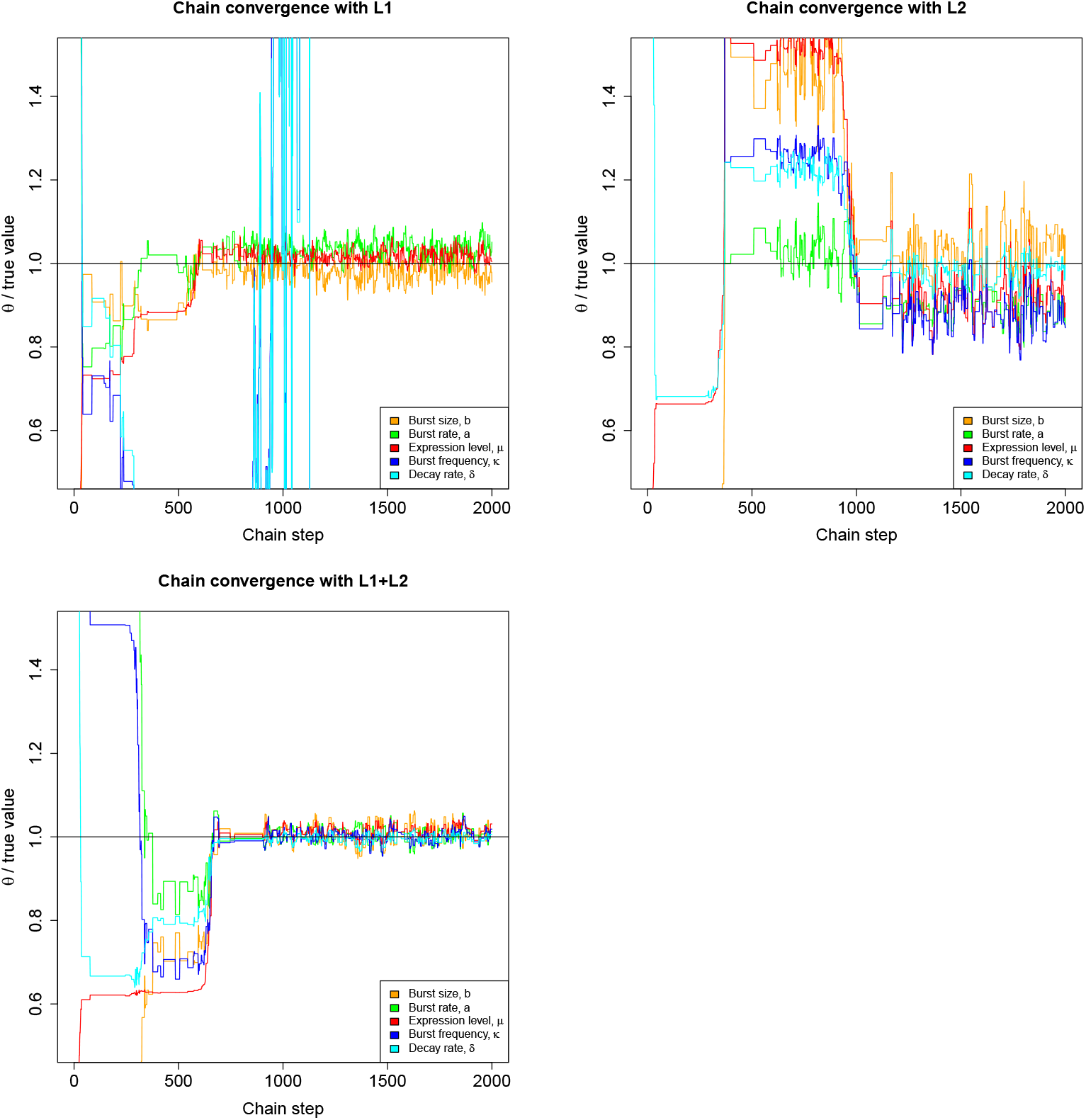
Convergence of Markov chains to true parameter values with simulated data for three different likelihood functions. The parameter values, *θ*, in the chain are divided by the true value to allow for joint visualisation, with the black horizontal line representing the target value.

The resulting posteriors (figure 4) indicate that the accuracy and precision of estimates for *a, b* and *μ* are improved by incorporating the single cell 4sU conversion data compared to relying solely on scRNA-seq data, which is because the cell-cell variance in the T>C rate is a function of the transcriptional noise (burstiness) of the gene as well as turnover and, therefore, including such information makes the estimation more robust. Likewise, we see that while conventional scRNA-seq may not resolve *κ* or *δ*, including the UMI count information with the conversion data also results in more precise and accurate estimates of these parameters. This is because the set of T>C conversions is a function of *a, b* and *δ*, while the UMI counts are a function of a and b. Therefore, including the UMI data improves inference of *a* and *b*, which reduces the error associated with *δ* in our joint inference approach.

**Figure 4:**
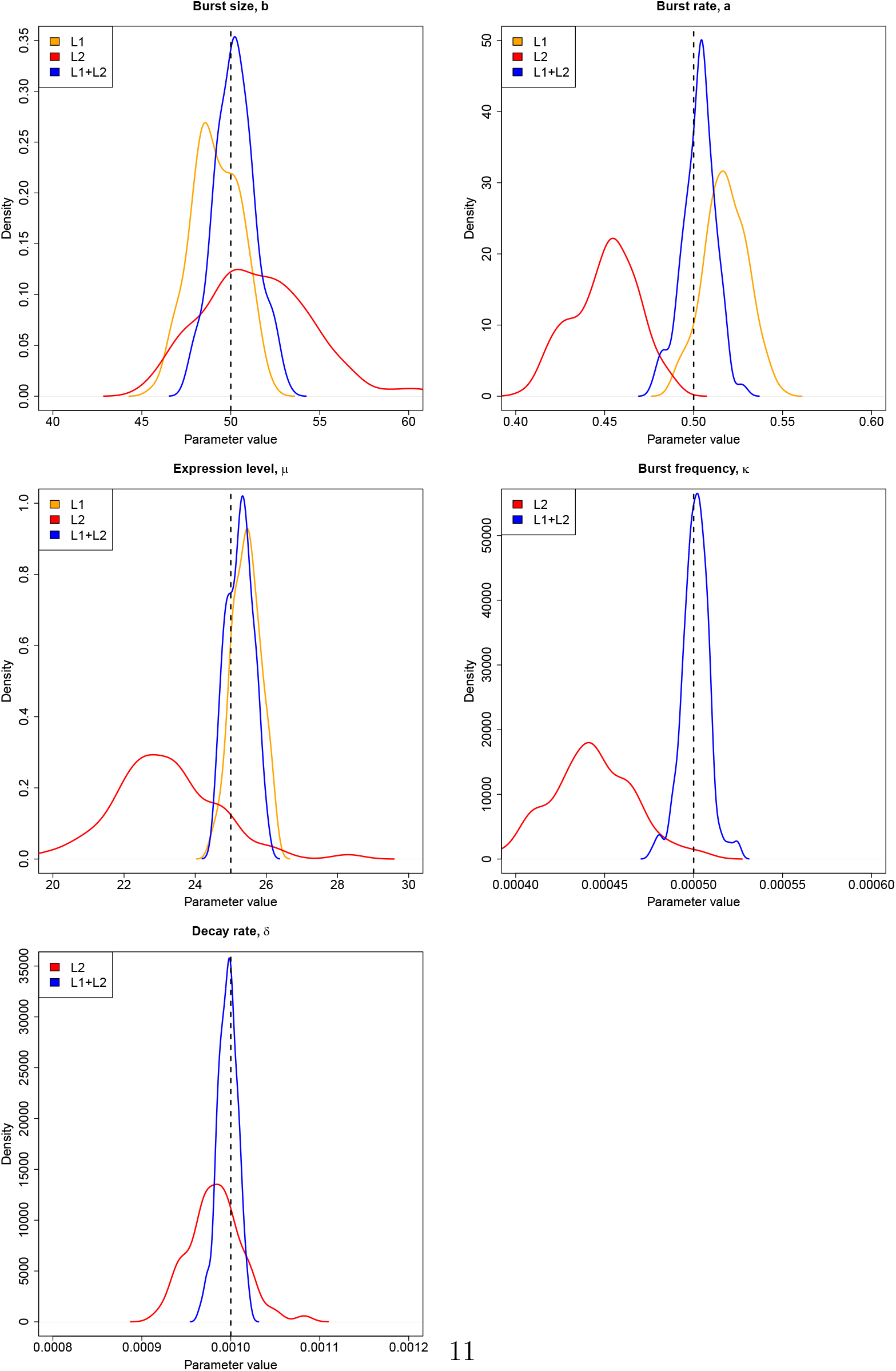
Probability density functions of each parameter derived from posteriors obtained using different likelihood functions, with the dashed black lines representing the true parameter values that were used to simulate the dataset upon which inference was carried out.

Overall, these results demonstrate the benefits a fully integrated analysis of time-resolved bursting dynamics using 4sU scRNA-seq data provides over more limited, separate analyses of subsets of the parameters by scRNA-seq (*a* and *b*) and bulk 4sU RNA-seq (*δ*). This is apparent in this example of a gene with moderate expression, high transcriptional noise and a transcript lifetime similar to the 4sU pulse duration.

### Inference on data from Qiu

We next tested our method on 4sU scRNA-seq data published in 2020 by Qiu et al, which used human K562 cells [38]. Inference on the data from Qiu was carried out for all genes with at least one read and observed T>C conversion in both the 4sU and control datasets, running the MCMC algorithm in parallel on each to obtain a posterior from model 1 and 2 (using two stages of inference, Methods). A number of these genes had to be discarded (Methods and SI), before selecting the final set of genes to be analysed which had sufficient confidence in all parameter estimates. Therefore, a maximum CV value of 0.365 was imposed for all parameter estimates obtained with both model 1 and 2 (ignoring *κ* and *δ* for model 1), so that only genes with no CV > 0.365 would be included. This makes use of the inference with model 1 as well as model 2 to maximise the robustness of our confidence quantification, leaving 606 genes as the final selected set.

For the selected genes we observe that the quality of our estimates depends upon the location of the gene within parameter space, as shown in figure 5, which depicts estimate vs CV for all parameters for stage 2 inference. It is clear that *CV*(*δ*) has an optimal (minimum) value for *δ* corresponding to an average transcript lifetime equal to the 4sU pulse duration (4 hours), with confidence decreasing bidirectionally. We also have increased confidence for genes with higher *μ* since estimates for such genes are informed by a greater volume of data. Likewise, genes with greater *b* have greater confidence because, firstly, increased *b* results in higher *μ*. Secondly, for a given *μ*, having a higher *b* implies lower *a*, meaning that the transcriptional noise is higher, resulting in a more heavily skewed transcript count distribution (across cells) which may be more precisely attributed to a region of parameter space. We do not see a visually obvious trend in confidence for *a*. This is because it is associated with higher expression level but lower transcriptional noise. Therefore, a gene with higher *a* has more data points with which to inform the estimate but a less skewed transcript count distribution, so that the effects on confidence tend to cancel each other out. The trend in confidence for *κ* is essentially dictated by the *a* and *δ* values for the gene.

**Figure 5:**
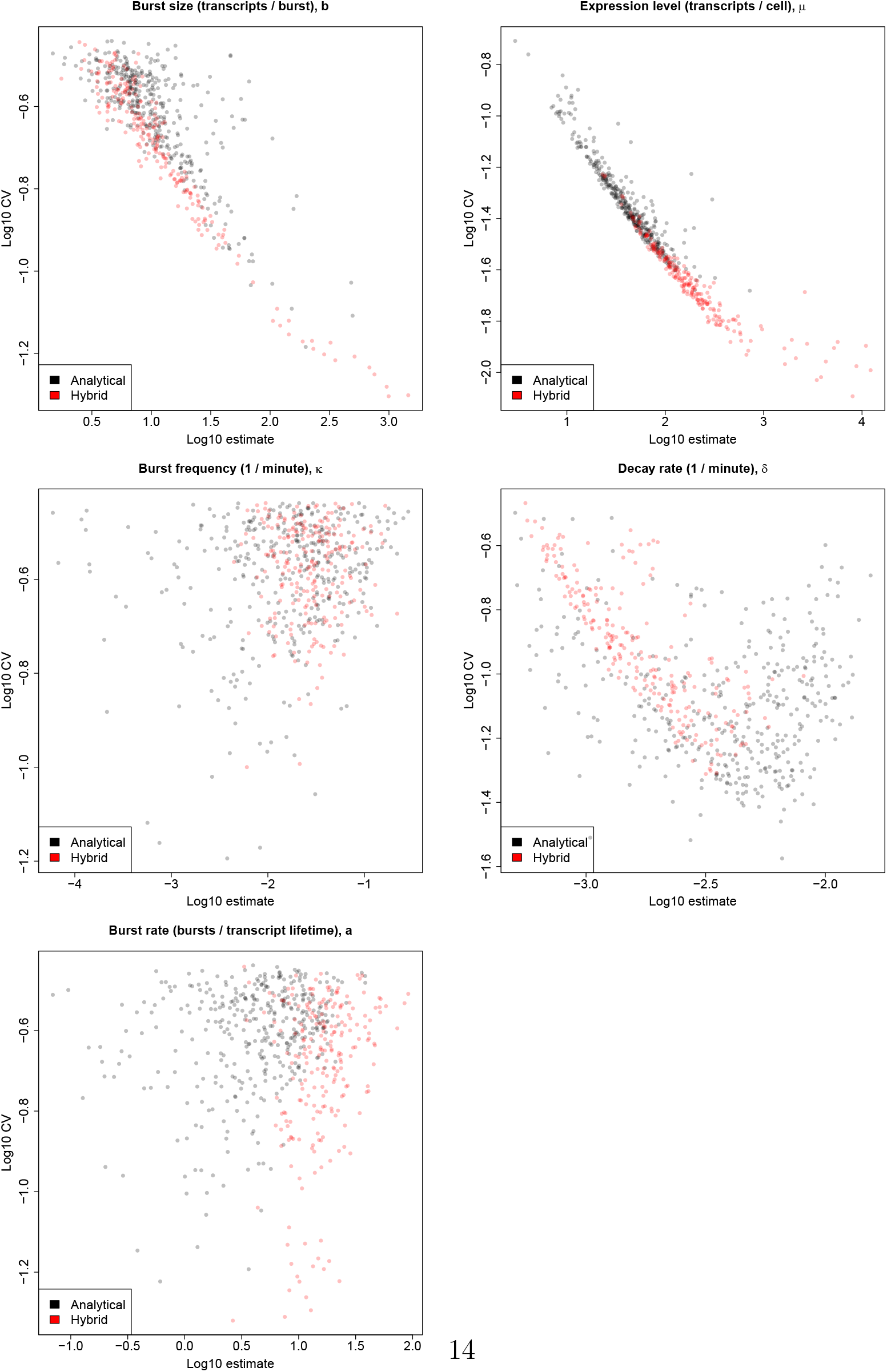
Estimates vs CVs of all parameters derived from model 2 posteriors for all 606 selected genes, with those obtained using the analytical and hybrid approaches displayed in black and red, respectively.

Instead of using a fully analytical likelihood to carry out inference with model 2, for some genes we must rely upon a hybrid approach which replaces part of the analytical likelihood function with a simulation-based boolean as with ABC (Methods). This occurs when genes lie within a region of parameter space such that the state space becomes too large or the solution to equation 8 becomes unstable. Figure 5 provides evidence supporting the reliability of the hybrid inference approach, since the genes for which this approach was used generally occupy the same regions of the plot as those treated with the analytical approach. This also illustrates the increased probability for a gene to reside within unstable parameter space, and therefore require the hybrid inference approach, when *μ* and *a* are higher and when *δ* is lower.

We reinforce our results by demonstrating a strong positive correlation about the diagonal between our estimates of *δ* and cell-matched values calculated in [36] for the same genes (SI, figure 15). Further assessment of our parameter estimation and confidence quantification was provided by carrying out inference on simulated data. This simulation-based validation differs from the previously described model comparison analysis (figures 3 and 4) in that experimental settings, such as cell number, cell capture efficiency and sequencing depth, were equivalent to those in the Qiu dataset rather than being idealised, and the bursting parameter values estimated for each of the 13236 genes we analysed were used as the true values for a corresponding simulated gene. Strong, tight correlations about the diagonal between estimates and true parameter values confirmed the capacity for the algorithm to recover known parameter values (SI).

Now that we have estimates for all parameters of interest, it is possible to demonstrate how the different aspects of the data feed into informing the joint probability of *θ* = (*a,b, μ, κ,δ*). Figure 6 illustrates some expected correlations, showing that *μ* correlates very strongly with the mean UMI count and that *δ* correlates very strongly with the 4sU - control T>C rate, since these values reflect the overall activity and turnover of the gene, respectively. We see that a correlates strongly against the CV of the UMI count, which reflects the relationship between bursting and cell-cell variability. It is also possible to demonstrate the aforementioned complex relationship between burstiness and the shape of the single-cell T>C count data, but not in a genome-wide manner since the effect is masked by variation in *μ* and *δ*. Therefore, we instead compare a pair of genes (*ATF5* and *TSN*) with very similar estimates for *μ* and *δ* but very different values of *a* (and therefore also *b* and *κ*), with *ATF5* being expressed in a far more bursty fashion. The estimates for the different parameters of these genes are given by table 1.

**Figure 6:**
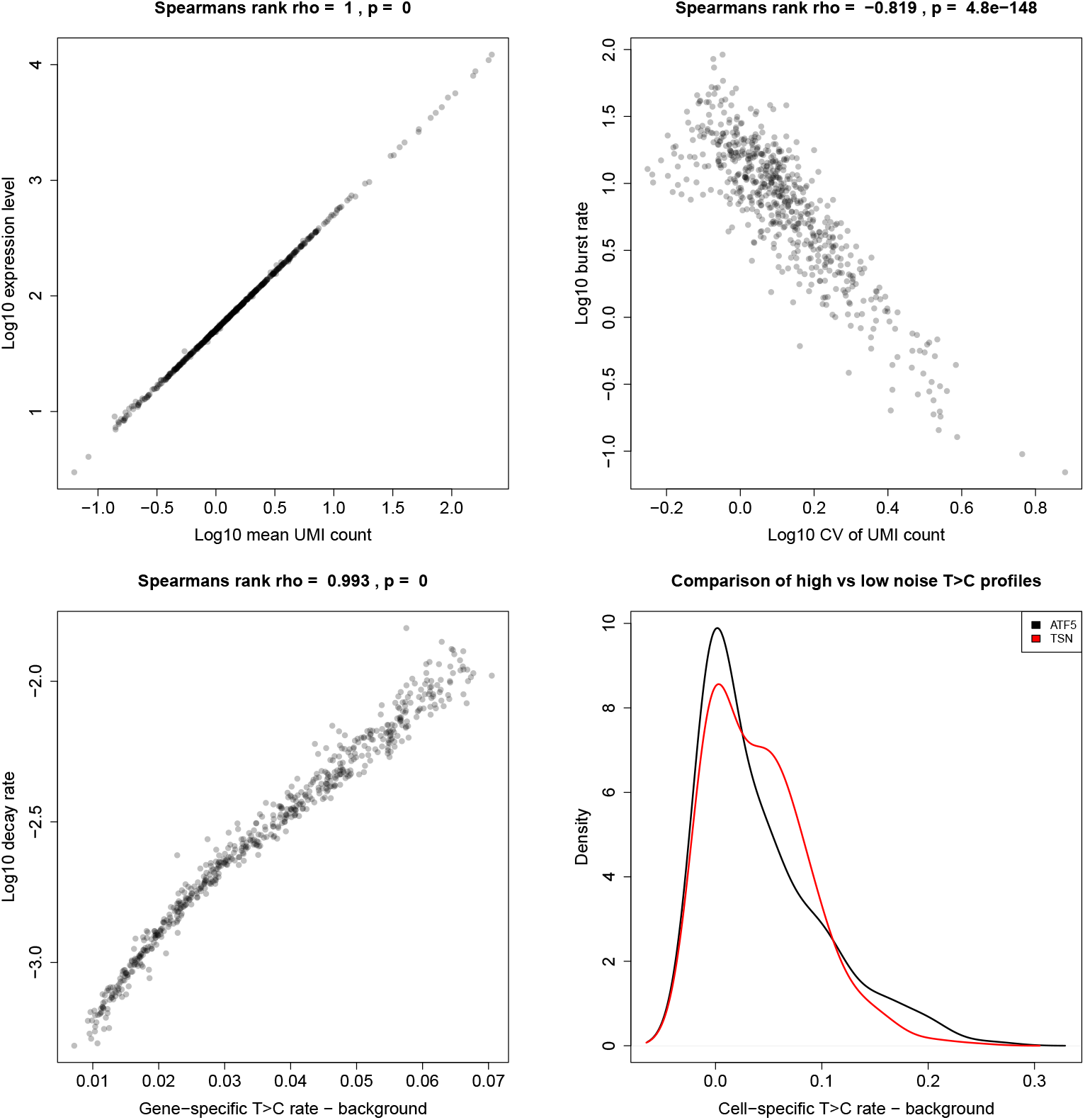
Correlations between statistics of the observable data and related bursting parameter estimates, with Spearman’s rank correlation strength (rho) and statistical significance (p) displayed. Bottom right compares the cell-specific T>C rates minus gene-specific background for the *ATF5* and *TSN* genes, which are expressed with high and low noise, respectively.

**Table 1:** Parameter estimates for the *ATF5* and *TSN* genes.

|  | ATF5 | TSN |
| --- | --- | --- |
| $b$ | 60.7 | 2.78 |
| $\mu$ | 60.2 | 58.3 |
| $\kappa$ | 0.00404 | 0.0873 |
| $\delta$ | 0.00403 | 0.00389 |
| $a$ | 1.00 | 22.4 |

The density plot in figure 6 compares the distribution of cell-specific T>C rates (minus gene-specific background) across all reads in the cell for the aforementioned pair of genes. There is a clear difference in the shape of the distribution, with the bursty gene having a greater density at either extreme while the gene with less noisy expression has a greater intermediate density. This is because large, infrequent bursting has a binarising effect, meaning that most cells either have a low or high T>C rate. Those with a low rate correspond to those which have had no bursts occur during the 4sU pulse, resulting in their entire transcript population comprising those surviving from before the pulse. Those with a high rate correspond to those which have had at least one burst occur during the pulse. Since the bursts tend to be large, this results in the majority of the transcript pool being comprised of newly synthesised transcripts. On the other hand, smaller, more frequent bursts causes the surviving transcripts to gradually become replaced by new transcripts in a more uniform manner across cells. Similarly to how scRNA-seq reveals differences in cell-cell variation in transcript counts for two genes with otherwise equal expression levels, 4sU scRNA-seq also reveals differences in cell-cell variation in new transcript proportions for two genes with otherwise equal decay rates.

Despite controlling for *μ* and *δ* in this pairwise comparison of a high vs low noise gene, the effect of bursting on cell-specific T>C rates shown in figure 6 is still somewhat obscured by the variable cell-specific capture efficiencies, *α*, present in the data. Therefore, datasets were simulated in the same manner as for the model comparison analysis, except λ_*s*_ = 0.001, and *α* =1 to totally control for the effect of capture efficiencies. Datasets were simulated for a gene with high noise and another with low noise with parameter values set as shown in table 2.

**Table 2:** Parameter values for simulated high and low noise genes.

|  | High noise | Low noise |
| --- | --- | --- |
| $b$ | 250 | 25 |
| $\mu$ | 250 | 250 |
| $\kappa$ | 0.001 | 0.01 |
| $\delta$ | 0.001 | 0.001 |
| $a$ | 1 | 10 |

The differential transition from surviving to new transcript pool for high and low noise genes is demonstrated in additional file 1, which shows the cell-specific T>C rate distributions for data simulated with different pulse durations. This illustrates the previously discussed effect of bursting on cell-cell turnover variation more clearly, visualising the bimodal vs unimodal transitions occurring under high vs low noise conditions with a video.

### Biological findings

Correlating the parameter estimates against each other for our 606 genes also reveals that genes with extremely high expression levels, the majority of which are mitochondrial genes, are able to achieve these high levels primarily by having very large bursts, rather than very frequent bursts or very stable transcripts, although the decay rates do appear somewhat constrained (figure 7). There may be biological upper limits on *κ* due to the various factors required to be in place to prime a gene for activity and, therefore, it may be preferable to instead increase active period duration (reduce *k_off_*), and therefore burst size, for very high expression levels [32]. A similar phenomenon has been observed previously, in which *MYC* overexpression lead to increased expression in target genes through increased burst duration and size, rather than increased burst frequency [45, 46]. Estimates for *κ* and *δ* are also positively correlated, perhaps because the cells are only able to tolerate a certain degree of noise in the expression of any given gene, otherwise too small a proportion may express the gene for the function to be fulfilled. This may manifest as a correlation between these two parameters to stabilise *a*, and thus the transcriptional noise.

**Figure 7:**
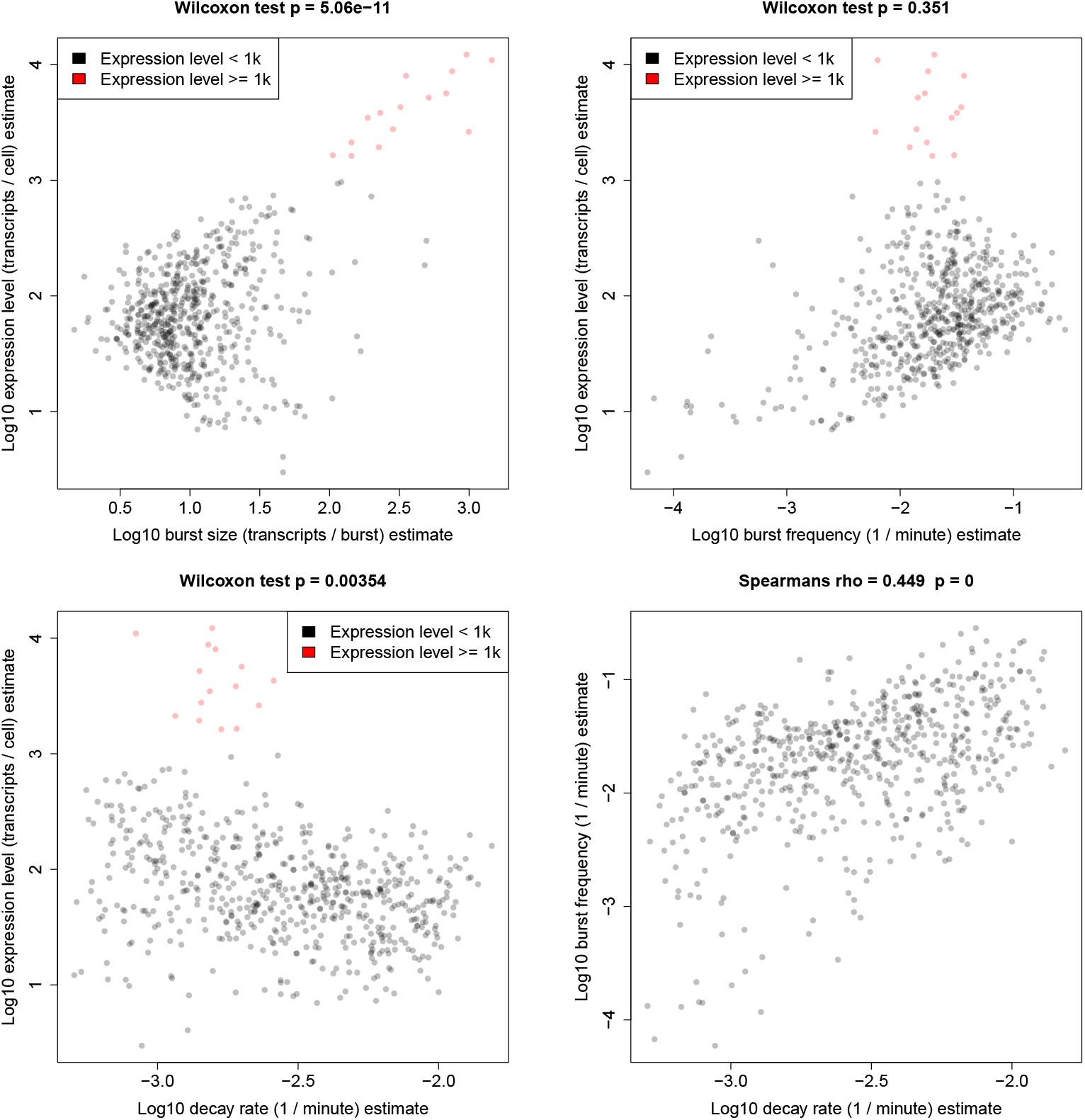
Estimates for different parameters plotted against each other. Statistical significance of difference in *b, κ* and *δ* for genes with very high (*μ* ≥ 1000) expression level vs other genes (*μ* < 1000) is shown with the p-value calculated using the Wilcoxon test. Also shown in the bottom right is the Spearman’s rank correlation strength (rho) and statistical significance (p) of *κ* against *δ*.

### Histone modifications and bursting

We next explored the relationship between HMs and transcriptional bursting dynamics with a metagene analysis carried out using ChIP-seq data for eight HMs (Methods). In this analysis, the mitochondrial genes were removed from our set of genes with high confidence parameter estimates, with 543 genes ultimately being included. Of the eight active HMs analysed, the profiles generally fall into the two previously described categeories [20], being either predominantly promoter-localised (H3K4me2, H3K4me3, H3K9ac, H3K27ac) or gene body (GB)-localised (H3K4me1, H3K36me3, H3K79me2, H4K20me1, SI). To better understand the association between HM profile and bursting parameters, the genes were split in half, sorted by parameter estimate for each of the five parameters. Metagene comparison reveals position-dependent associations for promoter-localised HMs, using H3K4me2 as an example (figure 8). It appears that HM presence at the promoter and through the GB is associated with increased *μ* and also *a*, while increased *κ* is specifically associated with promoter but not GB presence. Conversely, presence through the GB excluding the promoter region appears associated with increased *b* and reduced *δ*.

**Figure 8:**
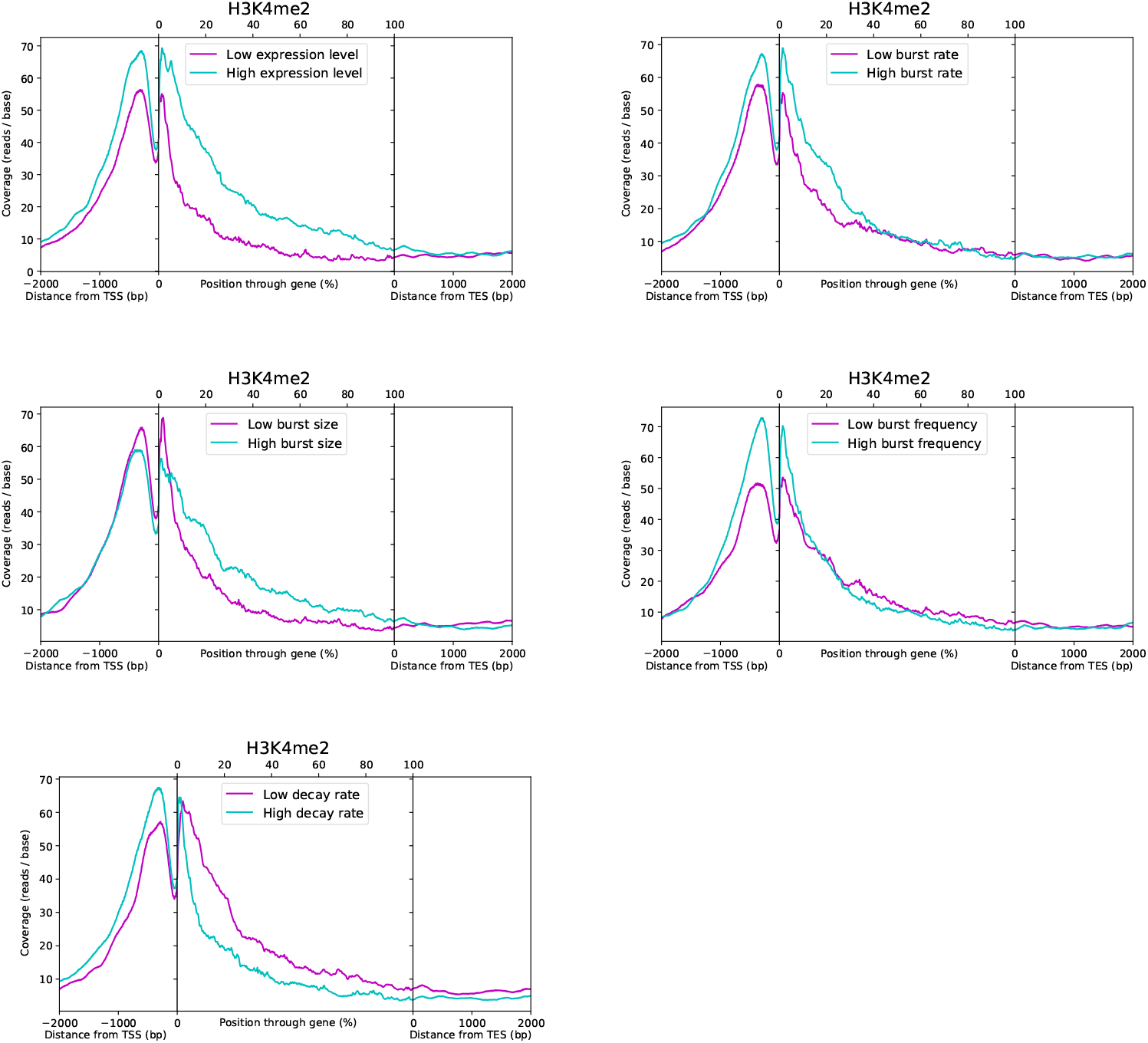
Metagene plots of H3k4me2 coverage, comparing profiles for the top and bottom 50% of genes when split according to their estimates for each parameter, denoted by high and low, as indicated.

This analysis builds upon a previous scRNA-seq study which correlated bursting parameter estimates with HM localisation by averaging the ChIP-seq coverage from 2000 bp upstream of the TSS to the TES for each gene [20]. They were unable to obtain estimates of *κ* or *δ* due to a lack of published data on transcript turnover rates for the cell type (hESCs). Our results are in agreement with [20] despite having a different cell type, but additional complexities are revealed which are only apparent with our metagene analysis combined with the capacity to estimate *κ* and *δ* afforded by 4sU scRNA-seq. For promoter-localised HMs, they report positive associations between HM presence and both *a* and *b*, whilst we demonstrate that the association with *b* is specific to the GB. We confirm that the association with *a* holds throughout both the promoter and GB, but show that this is a result of a promoter-specific positive *κ* association and a GB-specific negative *δ* association, thereby further demonstrating the advantages of 4sU scRNA-seq inference.

In order to statistically test these apparent associations, the average HM coverage values around the promoter and through the GB excluding the promoter were obtained for each HM (Methods), taking the average value from 2000 bp upstream of the TSS to 5% through the GB (−2000:5%) and from 5% through the gene body to the TES (5%:100%), respectively. Spearman’s rank correlation of the mean value for each promoter-localised HM against each parameter across our 543 genes confirmed the direction and quantified the strength (figure 9), as well as confirmed the statistical significance of the suspected associations (figure 10).

**Figure 9:**
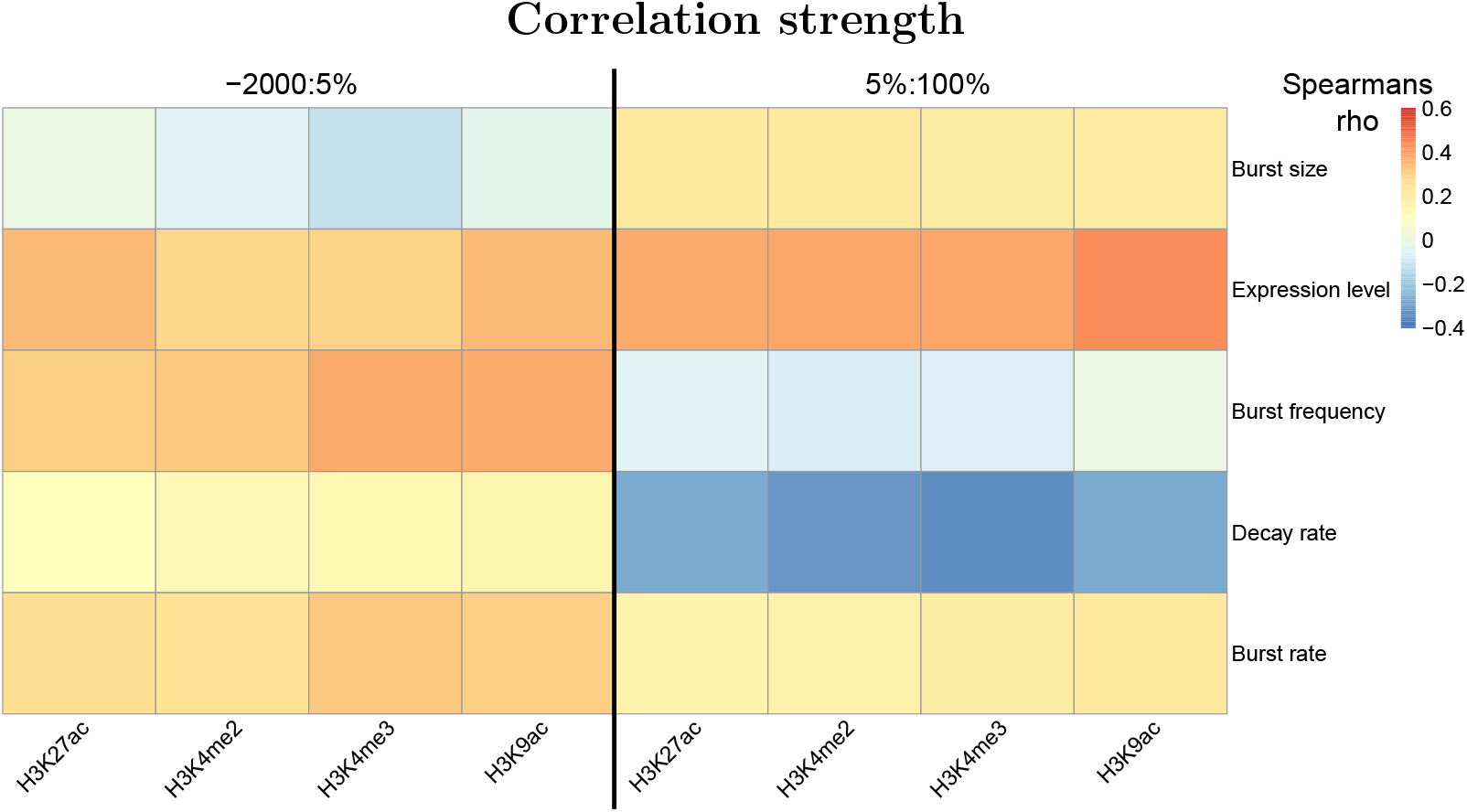
Heatmap showing the Spearman’s rank rho as the heat intensity value for the correlations between bursting parameter estimates and the mean promoter-localised HM coverage values across the −2000:5% and 5%:100% regions. More intense red or blue colouration indicates a stronger positive or negative correlation, respectively, while neutral indicates no/weak correlation.

**Figure 10:**
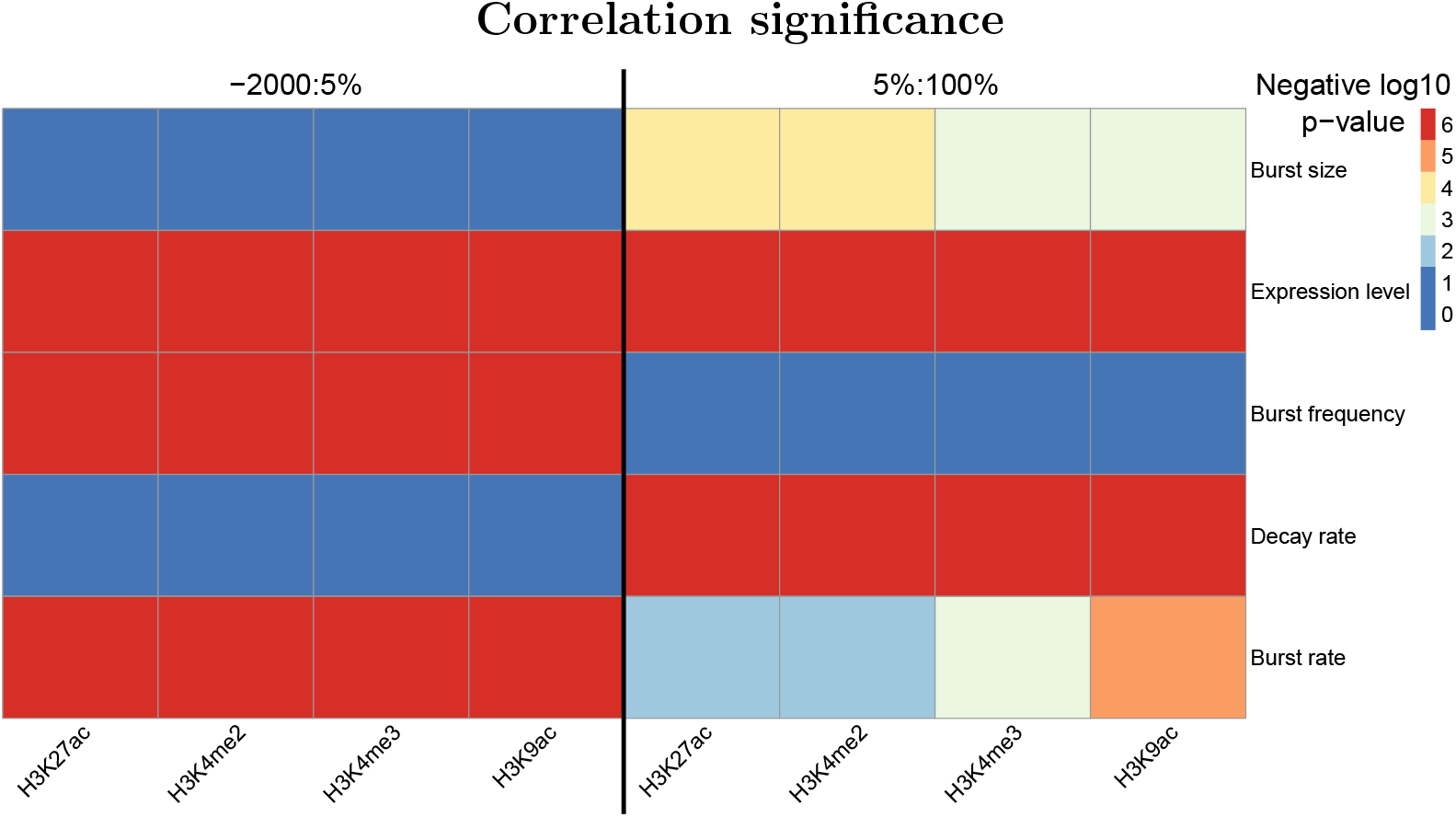
Heatmap showing the Spearman’s rank p-value (adjusted for multiple hypothesis testing) as the heat intensity value for the correlations between bursting parameter estimates and the mean promoter-localised HM coverage values across the −2000:5% and 5%:100% regions. The heat values are discretised, corresponding to negative log10 p-value thresholds. For example, the most intense blue indicates that, for the given correlation, 10^-2^ < *p*, meaning no statistical significance, the neutral colour indicates that 10^-4^ < *p* ≤ 10^-3^, while the most intense red indicates that *p* ≤ 10^-6^.

The association between promoter-localised HM presence and reduced decay rate is consistent with previous reports of a link between HMs and pre-RNA processing. The RNAP elongation speed may be modulated by HMs or they may be responsible for the recruitment of splicing factors [47, 48]. This could result in more stable RNA by ensuring correct splicing and/or polyadenylation. GB presence of promoter-localised HMs could also result in increased burst size by facilitating TSS-TES contact through the maintenance of the open chromatin state around the TES. Coupled with the free movement of RNAP through the GB, this may increase the burst size by allowing RNAPs to quickly and repeatedly generate multiple transcripts by promoting polymerase recycling [14].

## Discussion

With the inference approach presented here we demonstrate the capacity to obtain genome-wide estimates of the parameters governing transcriptional bursting dynamics and the timescales upon which they occur from a single dataset with no prior knowledge. By sampling from the full joint probability distributions of the parameter values given the data we are able to quantify confidence in our estimates and take into account the complex interdependencies between the different parameters and 4sU scRNA-seq data, revealing the regions of parameter space for which we have the most accurate and precise estimates. We show that the distribution of 4sU-induced T>C conversions across cells is shaped not only by the turnover rate and expression level of the gene, but also by the transcriptional noise, and that this information can therefore be used to improve estimates of burst rate (*a*) and burst size (b) beyond the level obtainable with conventional scRNA-seq. In this way, combining metabolic labelling and single cell resolution has an effect greater than the sum of their parts on inference power. Previous analysis of transcriptional bursting using 4sU scRNA-seq data has tapped into this idea by estimating the proportion of new transcripts (based on T>C conversions) in each cell for a particular gene and then using the standard deviation of this new to total ratio as a proxy for burstiness [43]. However, as clearly demonstrated by the video in additional file 1, this distribution, and therefore its standard deviation, is shaped not only by transcriptional noise but also by RNA turnover, and may be skewed by technical noise such as variation in capture efficiency. Therefore, along with the overall expression level, this needs to be explicitly accounted for in order to accurately quantify burstiness, as is naturally achieved with our mathematical model.

Having genome-wide estimates of the parameters governing transcriptional dynamics means that it is possible to use the variation which naturally exists between genes to examine the relationships between the different parameters and other features, such as HMs, instead of having to rely on experiments which artificially perturb the cells to gain insight via a single gene system. In agreement with previous reports [42], we find that the genes with very high expression levels are primarily mitochondrial genes. Going beyond this, we show that such activity levels are achieved by having large burst sizes rather than increased RNA stability or burst frequency, which we hypothesise could be due to biological constraints on the rate of switching between active and inactive states [32], potentially making it favourable to instead increase the duration of bursts, and therefore the burst size, as has similarly been observed for *MYC* -driven transcription [45, 46]. Whereas some studies have found the variation in decay rates (in mESCs) across genes to be an order of magnitude lower than for the other parameters, and therefore negligible [32], we found significant variation in K562 cells which was important to account for in order to properly estimate burst frequencies. Indeed, our analysis revealed an unexpected positive correlation between burst frequency and decay rate, resulting in the burst rate, and therefore transcriptional noise, being constrained. One may speculate that only noise levels within a certain range are tolerated, with extreme values resulting in too few cells expressing the gene for a given function to be achieved, such as the appropriate proportion of cells in an isogenic population undergoing differentiation [11, 49], manifesting as the observed correlation.

Examining the relationship between bursting parameters and HMs genomewide produced results consistent with but advancing upon previous work [20]. Combining our metagene analysis with the additional information provided by 4sU scRNA-seq over inference on conventional scRNA-seq reveals intricacies that were not previously apparent. The presence of GB-localised HMs throughout the gene is generally associated with increased burst rate (bursts per transcript lifetime) via increased burst frequency (bursts per minute), while promoter-localised HMs are only associated with increased burst frequency when found around the TSS. Their presence further downstream remains associated with increased burst rate, and therefore reduced transcriptional noise, but through reduced decay rate rather than increased burst frequency. The association with reduced decay rate may be related to the previously documented influence of HMs on pre-RNA processing, which is achieved, for example, by modulating RNAP elongation speed and/or by recruiting splicing factors [47, 48]. This may increase RNA stability by reducing the probability of incorrect splicing or polyadenylation. Presence of promoter-localised HMs throughout the GB but not at the TSS is also associated with increased burst size. Downstream presence could facilitate interactions between the TSS and the TES by maintaining the open chromatin state around the TES. This, along with maintaining the free movement of RNAP through the GB, could promote polymerase recycling and therefore increased burst size by allowing RNAPs to quickly and repeatedly fire off multiple transcripts during an active period [14].

The inference approach described here is generally applicable to 4sU scRNA-seq datasets which have RNA spike-ins and UMIs for any organism or cell type. Furthermore, the model could easily be expanded to integrate an arbitrarily large number of repeat experiments by extending the Markov chain according to the product of the likelihood functions of each dataset. Indeed, such a scheme which utilised datasets with different 4sU pulse durations could theoretically characterise the transcriptional dynamics of all genes genome-wide. For example, inference carried out using two datasets with long and short pulse durations would facilitate estimates for genes with long and short transcript lifetimes, respectively. The implementation of our alternative, ABC-like inference approach could also be useful in future for handling more complex, analytically intractable mathematical models. A caveat of our analysis is the asynchronisation of the cell cycle phase across the population. This may confound the results in two ways, firstly because different phases have a different cellular environment, influencing the global transcriptional dynamics and causing variation in the underlying parameter values for the same gene between cells in different phases. Secondly, there is variation in the copy number of genes throughout the cell cycle, with an unknown proportion of cells having one or two copies of each nuclear gene. Confounding effects on the inference could be resolved by separation of the different subpopulations of cells by cell cycle phase using, for example, fluorescence-activated cell sorting prior to sequencing [33], and/or by using allele-specific/sensitive scRNA-seq approaches combined with metabolic labelling [17, 34]. As 4sU scRNA-seq data becomes more common place and there are improvements in capture efficiencies, sequencing depths and cell numbers, it will be possible to robustly infer time-resolved transcriptional bursting dynamics for a far greater number of genes from a single experimental set up. Our findings on burst dynamics and their associations with HMs could be a valuable starting point to inform future experimental work investigating this area, while further application of our method beyond what is presented here might hint at other, novel mechanistic relations.

## Conclusions

In conclusion, we have developed a mathematical model to maximally exploit the power of 4sU scRNA-seq datasets to examine transcriptional bursting, tapping into the synergy between single-cell resolution and 4sU labelling which manifests in the cell-specific T>C rate distributions. The advantages over conventional scRNA-seq were demonstrated in detail using small-scale simulations and performance of the algorithm across parameter space was validated with large-scale simulations. We applied our inference approach to published 4sU scRNA-seq data to obtain genome-wide joint parameter estimates and confidence quantifications, finding an unexpected correlation between burst frequency and decay rate, and that genes with extremely high expression levels achieve this primarily through increased burst size. Finally, we linked our estimates with published ChIP-seq data, revealing position-dependent associations between different histone modifications and parameter estimates which only become apparent with 4sU scRNA-seq as opposed to conventional scRNA-seq.

## Methods

### Data processing and analysis

#### 4sU scRNA-seq

The main datasets that were used for parameter inference in this study were produced in Qiu et al 2020 [38], downloaded from the GEO series GSE141851. Two datasets from this series were used, both using K562 cells; a negative control dataset with TFEA chemical conversion treatment but with no 4sU added, and another dataset which had 4sU added 4 hours before chemical treatment, with GEO sample IDs GSM4512696 and GSM4512697, respectively. These are Drop-seq datasets and thus were processed according to the “Drop-seq alignment cookbook” (https://mccarrolllab.org/wp-content/uploads/2016/03/Drop-seqAlignmentCookbookv1.2Jan2016.pdf). A custom Python script was used to carry out trimming of read pairs with any base with phred quality ≤ 10, and to clip adaptor and polyA tail sequences. The trimmed reads were then aligned to the primary human genome assembly (GRCh38.p13), the fasta file for which was obtained from gencode (https://www.gencodegenes.org/human/), using bwa to build the genome index and for the actual alignment [50]. Custom Python scripts were then used to map the aligned reads with mapq score ≥ 10 to their genes according to the gencode.v36 primary human genome assembly annotation gtf file, before extracting cell-specific (using the cell ID part of the read 1 barcode) UMI counts and total read counts for each gene, along with gene-specific, cell-specific information for each read about the number of genomic T bases (found in the fasta sequence across the aligned read positions) and the number of those which were converted to C bases in the read sequence. Cell selection was then carried out to exclude those cell IDs corresponding to empty droplets by ordering the cell IDs based on the total number of corresponding read pairs and then selecting the top 400 or 795 IDs for the control and 4sU dataset, respectively, as specified in [38]. The control dataset was then used to derive the gene-specific background T>C conversion rates, *λ_s_*, based on the proportion of genomic Ts which were converted to Cs across all reads across all selected cells for the given gene.

#### ChIP-seq

Publicly available ChIP-seq datasets for eight active HMs produced with K562 cells were downloaded for our analysis. A H3K4me3 ChIP-seq dataset was obtained from the GEO series GSE108323 with sample ID GSM2895356, which had been processed with alignment to the hg19 human genome build [51]. Seven more ChIP-seq datasets, which had also been processed with alignment to the hg19 human genome build, were obtained from the GEO series GSE29611 with sample IDs GSM733651, GSM733653, GSM733656, GSM733675, GSM733692 and GSM733714, corresponding to H3K4me2, H3K79me2, H3K27ac, H4K20me1, H3K4me1 and H3K36me3, respectively [52]. The position and read count information from these datasets was used to obtain the single-base resolution coverage values for each HM. These values were associated with their corresponding genes using the information from the comprehensive gene annotation hg19 gtf downloaded from Gencode. Analysis of the correlations between bursting parameter estimates and HM coverage at different sections of the gene was carried out by taking the average coverage value for all bases across the specified section (e.g. from 2k bp upstream of the TSS to the TES) for each gene, so a single value is obtained per gene per HM. Metagene plots were produced by averaging the coverage values for each position through/around the gene across all specified genes, similarly to the metagene analysis described in [53].

#### Mathematical modelling

In general, we model bursty transcription as a stochastic process closely related to the standard two-state model, as many previous works have [9, 10, 14]. The two-state model has four possible processes of gene activation, gene repression, transcription and degradation, where transcription may only occur with the gene in an active state while degradation acts continuously. This is represented by the following chemical reaction scheme

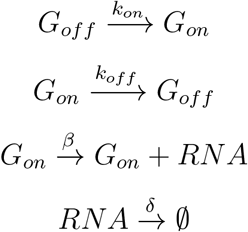

in which *k_on_, k_off_, β* and *δ* represent the rate constants for gene activation, gene repression, transcription and RNA degradation, respectively, while *G_off_*, *G_on_* and *RNA* represent the different species of repressed gene, active gene and transcript, respectively. A schematic representation of the system is shown in figure 11.

**Figure 11:**
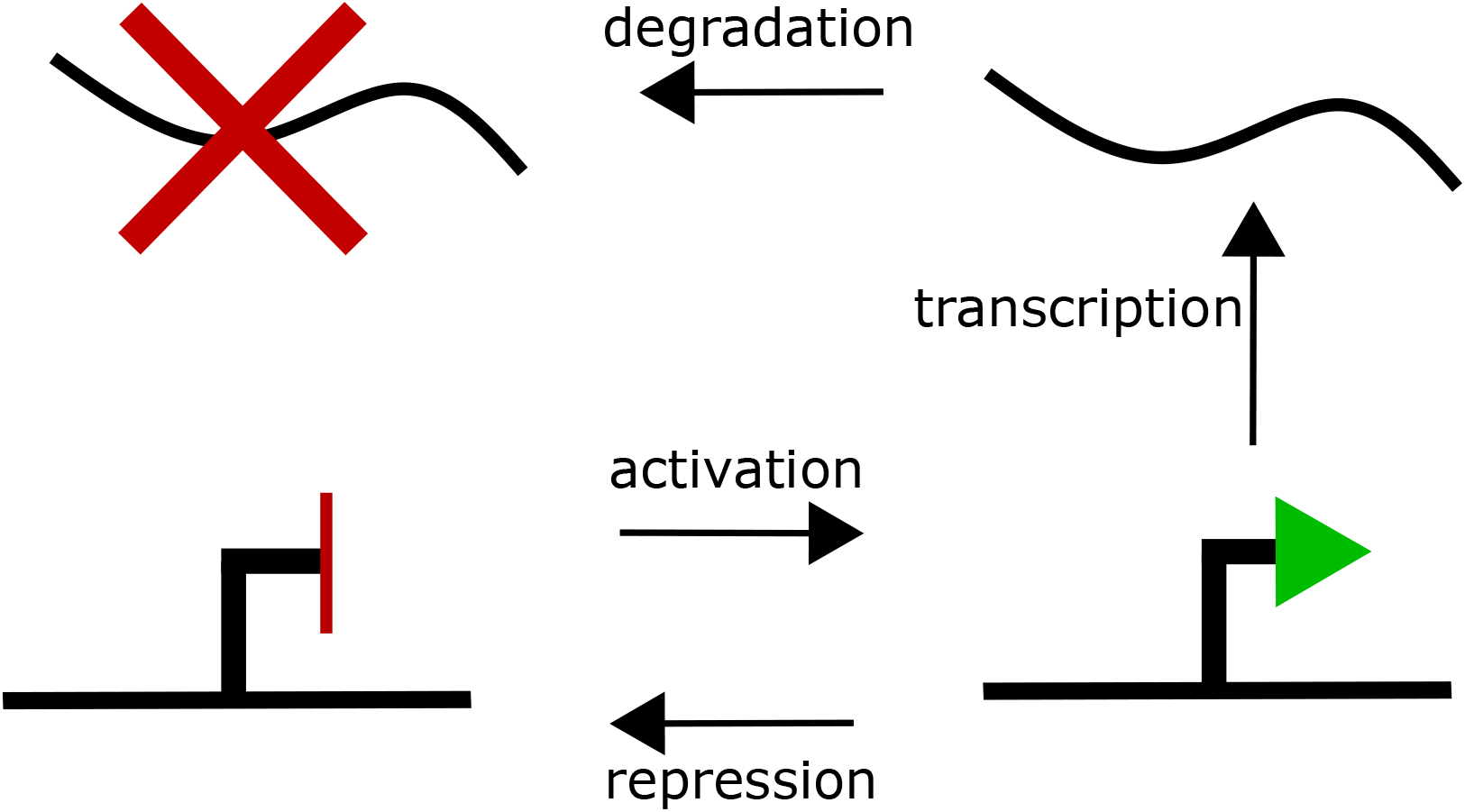
Schematic representation of the two state model, with the four reactions (activation, repression, transcription and degradation) acting on the three species (repressed gene, active gene and transcript).

With this model we have burst frequency, 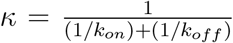 and burst size, 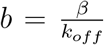, and we recall the burst rate, 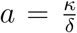. Aiming to understand bursting and its timescales specifically, we make the assumption that bursts occur instantaneously, arrive according to a poisson process and burst in a geometric fashion, which is valid when *δ* << *k_off_* since a transcript produced in a given burst is unlikely to have degraded before the burst is over [7, 54], and when *k_on_* << *k_off_*, which is supported by the parameter estimates reported in [32]. This model simplifies 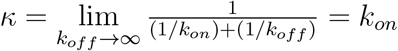 while *b* remains finite with 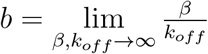 [55].

#### Model 1

The first model aims to model the observed unique molecular identifier (UMI) counts of a given cell, *l*, from the estimated capture efficiency (SI) of that cell, *α*, in a similar fashion to the technical noise model outlined in [56]. The capture efficiency, *α*, represents the transcript detection rate for that cell (probability of at least one read corresponding to a particular transcript). Based on the the instantaneous bursting version of the two-state model described above, the steady state distribution of the transcript count, *m*, can be derived directly from the master equation and corresponds to the negative binomial distribution [10, 54, 57, 55]

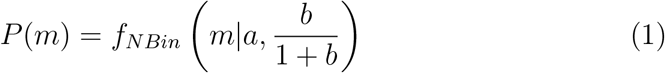

where

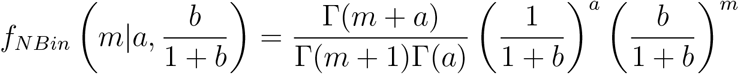

The full derivation is available in [55]. We may then model the probability distribution of observing *l* UMIs given *m* transcripts in the cell with a capture efficiency of *α*, as a poisson approximation of the true binomial process

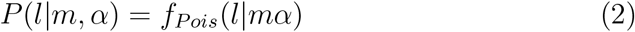

where

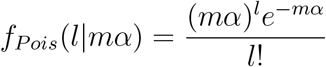

which is valid when *α* is small. We model the observed data, linked by the unobserved steady state transcript distribution by compounding equations 1 and 2 across the state space of *m* and marginalise

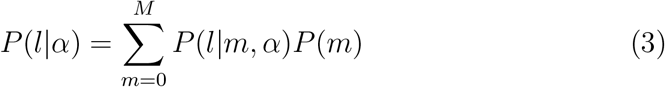

where *M* is an upper bound corresponding to the 0.9999 quantile of equation 1, which avoids summing to ∞, achieving a finite state projection (FSP) [58, 59] with an error of 0.0001. This leads us to the likelihood function of model 1 by taking the product of equation 3 across all cells in the data

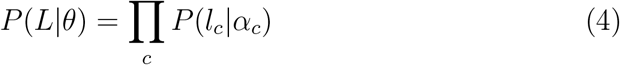

where *l_c_* and *α_c_* represent the observed UMI count (for the given gene) and capture efficiency for cell *c*, respectively, and *L* = (*l_x_*,…, *l_k_*), with *k* cells in total in the data and *θ* = (*a, b, δ*). Since we wish to infer the values of *θ* for each gene from the data using this model, we aim to obtain the posterior

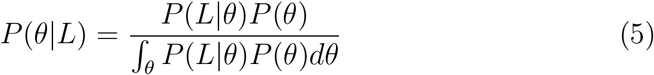

which we achieve through MCMC sampling.

#### Model 2

We will now construct a model which unifies the UMI and T>C conversion aspects of the data with the aim of understanding both bursting dynamics and the timescale upon which they occur. First of all we define *τ* = *tδ* where t is the time before sequencing at which the 4sU nucleotides were added to the cells, otherwise known as the pulse duration. *τ* therefore represents unitless time in terms of transcript lifetimes. Next we must obtain the probability mass function of the number of transcripts surviving to the sequencing point which were produced before the 4sU was added, otherwise known as the surviving transcripts, *s*. This distribution, *P*(*s*), may be understood as the time-decay of the steady state distribution, *P*(*m*), where we have 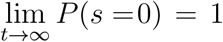 and *P*(*s*|*t* = 0) = *P*(*m*) when *δ* > 0. Degradation acts upon each individual transcript molecule with rate *δ*, and therefore the probability of a given transcript produced before 4sU was added surviving is 1 – *F_Exp_*(*X* ≤ *t*|*δ*) = *f_Pois_*(0|*τ*). Therefore, the probability of having s transcripts surviving given *m* originally is

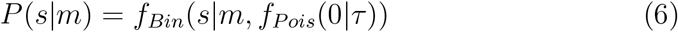

where

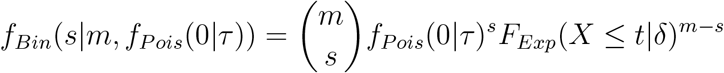

and

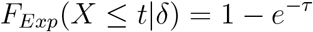

giving the conditional distribution of s. Compounding this with the steady state distribution (equation 1) we obtain the marginal

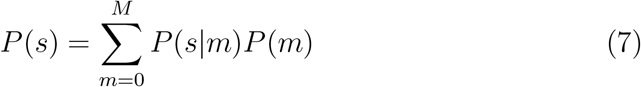

where *M* is an upper bound corresponding to the 0.99999 quantile of equation 1 for a FSP with error 0.00001. Next we obtain the probability mass function of the newly synthesised transcript count, *P*(*n*), for those transcripts that were produced after the 4sU was added and therefore have a higher T>C conversion rate than the background. This may be understood in reverse to *P*(*s*), as it describes the convergence of the newly synthesised transcript count from a point mass at zero to the steady state distribution where we have *P*(*n* = 0|*t* = 0) = 1 and 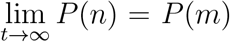 when *a,b,δ* > 0. An approximate solution to such a distribution was derived as a model of translation in [54] though the assumed relationships apply here. The solution is

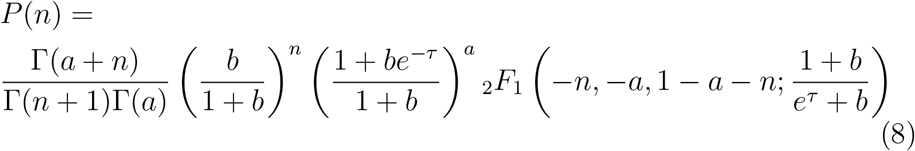

which is valid when *k_off_* >> *δ* and *τ* >> *δ/k_off_*, where _2_*F*_1_ refers to the hypergeometric function. Next, we obtain the probability distribution of transcripts at steady state conditional on our observed cell-specific capture efficiency, *α*, and UMI count, *l*, by using equations 1 and 2

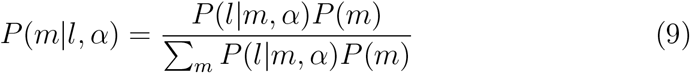

Now we describe the probability distribution of *n* conditional on *m* as the joint distribution of *n* and *s*

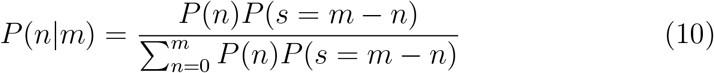

with the convolution 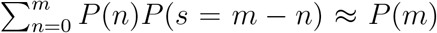 being used as a normalising value in place of *P*(*m*) due to the approximate nature of *P*(*n*), ensuring that 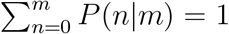. It is now possible to model the number of T>C conversions observed in a given read conditional on *m*, where we have expanded and built upon the poisson mixture model of conversions described in [36] and compounding with equation 10

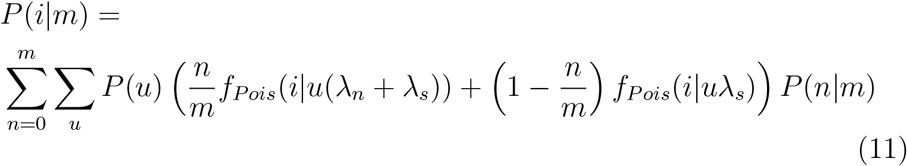

where *P*(*u*) is the gene-specific empirical probability mass function of observing *u* uracils across the fasta sequence corresponding to a given read’s mapping position. λ_*s*_ is the gene-specific background conversion rate observed in the control dataset (without the chemical conversion step) which represents conversion due to random mutations or other sources outside of chemical conversion. λ_*n*_ is the gene-invariant conversion rate due to 4sU incorporation and conversion which was estimated from the data (SI). We may now model the cell-specific T>C conversion rate for the given gene by compounding equations 9 and 11

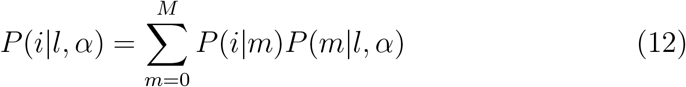

where *M* is an upper bound corresponding to the 0.9999 quantile of equation 1, again giving a FSP with error 0.0001. We are finally in a position to complete the model and link all our observables together. The observed counts of conversions in each cell may be represented by *y*, where *y_i_* is the number of reads that have i conversions. Therefore, the cell-specific observed distribution of conversions per read may be understood as a multinomial distribution with a probability vector determined by equation 12

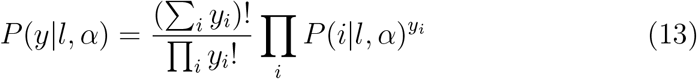

enabling us to model the conversion data conditional on the UMI data. A likelihood function may now be obtained with

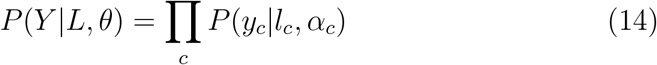

where *y_c_* is the conversions per read distribution observed in cell *c* and *Y* = (*y*_1_,…,*y_k_*) where *y_c,i_* is the number of reads with *i* conversions in cell *c* for the given gene. The final, full likelihood function is now defined as the product of equations 4 and 14

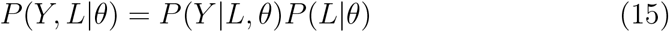

As in equation 5, MCMC sampling was used to obtain

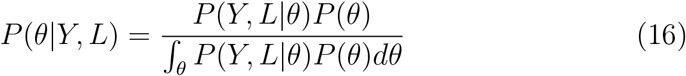

One thing to note about model 2 is that equation 8 is an approximate solution and breaks down in certain regions of parameter space. When *a* and/or *b* become too large and/or *τ* becomes too small, the function will oscillate around the true probability distribution function, with these oscillations quickly becoming more extreme to the point that the approximate solution gives negative probability values. The solution can be said to become unstable in these regions of parameter space, and therefore such regions will be referred to as unstable parameter space. If a gene is found to reside within an unstable region of parameter space then an alternative to the fully analytical likelihood function must be used. This alternative is also used when the state space of a gene is high enough to require an unfeasibly large number of calculations to compute the model 2 likelihood, which is only the case for a small handful of genes with extremely high expression levels.

### Markov chain Monte Carlo algorithm

#### Metropolis-Hastings algorithm

MCMC was employed in order to sample from the posterior distributions outlined in equations 5 or 16 using the Metropolis-Hastings algorithm [60], where *P*(*θ*) represents the prior distribution, which in this case is defined to be a multivariate uniform distribution. The chain is randomly initialised, *θ*_1_, where *P*(*X*|*θ*_1_)*P*(*θ*_1_) > 0 and *X* is the dataset. At each step, *j*, in the chain, a new proposal, *θ*\*, is drawn from the proposal distribution, *ρ*(*θ*\*|*θ_j_*). This is accepted with a probability given by the likelihood function

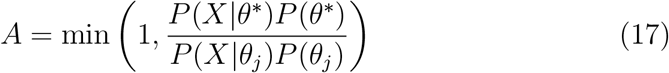

If the proposal is accepted then *θ*_*j*+1_ = *θ*\*, otherwise *θ*_*j*+1_ = *θ_j_*, and the process repeats until 10000 steps have been completed. Note that the probably intractable integrals in the denominators of equations 5 and 16 cancel out to allow the acceptance probability the be calculated with only the likelihood function and the prior distribution. In our special case with uniform priors, these also cancel out, only serving to reject proposals outside of the plausible ranges of parameter space as defined by the priors. Therefore, the Markov chain converges to the posterior distribution according to the gradient of the likelihood function. Posterior distributions were produced from the sampled chain using the last 7501 steps with a thinning factor of 5, where only every 5th point in the chain is used in order to reduce dependency between points, resulting in smoother probability density functions and a sample size of 1501.

#### Adaptive proposal distributions

Adaptive proposal distributions were used to optimise chain mixing flexibly for inference on large numbers of genes with different posterior distributions while minimising the user input and testing required. The chains were run for a total of 10000 steps, with initial steps using a multivariate normal proposal distribution, 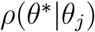, which is modified according to the Adaptive Scaling Metropolis algorithm described in [61], where Σ is the covariance matrix with zeros in every position except the diagonal, where *diag*(Σ_*j*_) = *θ_j_*/10. This scales the jump size (proposal standard deviations), aiming to eventually result in an overall acceptance rate (proportion of accepted proposals) which is considered optimal (0.234). However, this adaptive scaling modifies each element of the diagonal according to a single scalar value and does allow for covariances between different parameters in the proposal. Therefore, after the 10th accepted proposal, the proposal distribution switches to a multivariate adaptivity approach with a covariance matrix, Σ, that aims to approximate the covariance matrix of the posterior distribution, which is the adaptive Metropolis approach of [62]. Σ is calculated as described in [63], although only steps [max(1, *j* – 250), *j*] of the chain are used rather than the full history of the chain in order to preserve a quasi-memorylessness property

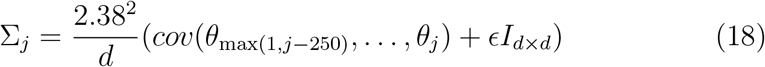

where *ϵ* =10^-10^, *I_d×d_* is the *d* by *d* dimensional identity matrix and *d* = 3 because *θ* = (*α,b,δ*). Then the proposal may be drawn as 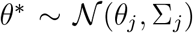 where

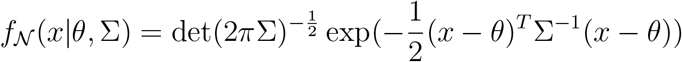

This covariance-based approach is not used until at least a few proposals have been accepted in order to avoid numerical problems when calculating the covariance matrix of the posterior.

#### MCMC with analytical/numerical hybrid likelihood

As previously mentioned, when carrying out inference using model 2, calculating equation 15 may be prohibited by the gene residing in an unstable region of parameter space or by the state space being too large. In this case, instead of calculating the likelihood function analytically, an analytical-numerical hybrid may be used. Equation 4 from model 1 represents the analytical portion, while the T>C conversion data is assessed using an approximate Bayesian computation (ABC) approach, as outlined in [64]. Under the usual scheme, for model 2 the proposal will be accepted according to

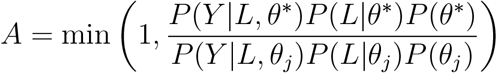

Under the hybrid scheme, this becomes

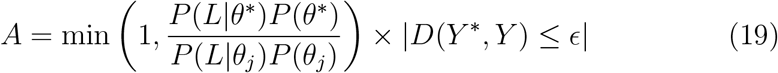

where the value of the right-most term is 1 or 0 depending on whether the condition is met or not, respectively. Here, *Y*\* represents a simulated dataset of T>C conversions produced with *θ*\*, *D*(*Y*\*, *Y*) represents a measure of the distance between the simulated T>C conversion data and that observed in the real data and e is the maximum distance value, above which the proposal will be rejected. *Y*\* is generated by drawing the steady state transcript counts for *N* cells at the start of the pulse

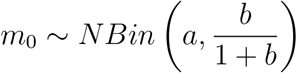

the surviving transcript counts at the end of the pulse

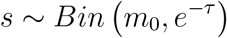

the number of bursts occurring during the pulse

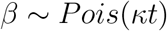

the times during the pulse at which each burst occurs

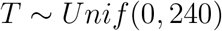

the size of each burst

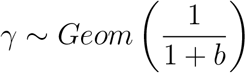

where

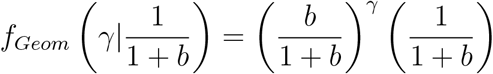

and the number of newly synthesised transcripts from each burst which survive to the end of the pulse

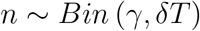

Then the total transcripts in each simulated cell at the end of the pulse is 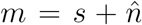, summing the number of transcripts surviving from each burst occurring during the pulse as 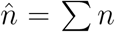. The total number of UMIs in each cell is then drawn as

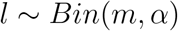

where *α* is randomly sampled without replacement from our list of capture efficiency estimates (SI). The list is repeated depending on the number of cells used to simulate the dataset, e.g. if 7950 cells are used the list is repeated 10 times since there are 795 cells and thus values of *α*. Now neglecting the single cell aspect of the data, we draw the total number of reads across all cells as

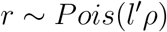

where *l*’ = Σ*l* is the sum of the UMI counts across all cells and *ρ* is the average number of reads per UMI observed for the given gene in the real data. Each read is assigned as corresponding to a newly synthesised or surviving transcript with probability *n*’/(*n*’ + *s*’) and *s*’/(*n*’ + *s*’), respectively, where 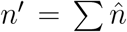 and *s*’ = Σ *s* represent the total new and surviving transcripts across all cells, respectively. We then randomly assign the number of uracils, *u*, that each read has by sampling from the gene-specific empirical distribution of uracils per read observed in the real data, *P*(*u*). Finally, the number of conversions in each read is sampled for new reads as

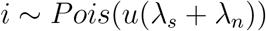

or for surviving reads as

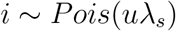

where λ_*s*_ and λ_*n*_ refer, as previously mentioned, to the gene-specific background conversion rate obtained from the control dataset and to the estimated gene-invariant conversion rate due to 4sU incorporation and chemical conversion (SI). From this we have our simulated dataset, *Y*\*, where 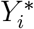 represents the total number of reads with *i* conversions across all simulated cells. We then take the real dataset, *Y*, again neglecting the single cell aspect so that *Y_i_* represents the total number of reads with *i* conversions observed across all cells. The distance metric between the two datasets is calculated based on the two cumulative distribution functions

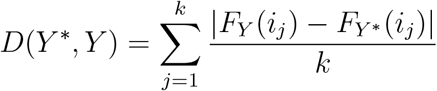

with *i* = (*i*_1_,…, *i_k_*), giving the mean of the absolute value of the difference between the cumulative distribution functions of the real and simulated data across all *i* observed in *Y*. This is substituted into equation 19 to complete the hybrid MCMC scheme.

#### Two-stage inference approach

A two stage approach for inference of *θ* was utilised, implemented via custom R and Python scripts. This was carried out initially with model 1, using equation 4 as the likelihood function to analytically obtain the posterior distribution, *P*(*θ*|*L*), of each gene. This is equivalent to treating the dataset as scRNA-seq data, ignoring the time-variant information of T>Cs provided by the 4sU aspect. An empirical estimate of the mean expression, μ, is calculated as

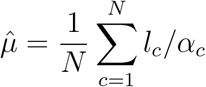

where *N* = 795 is the number of cells in the dataset. The chains were initialised by drawing log-uniform random variables

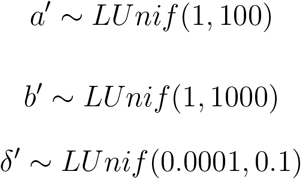

where

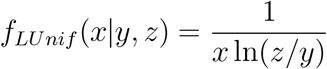

with support [*y, z*] for *y* > 0. Then set 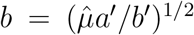, *a* = *ba*’/*b*’ and *δ* = *δ*’ to initialise *θ*_1_ = (*a,b, δ*), as long as *P*(*θ*|*L*) > 0, otherwise re-draw. The chain then proceeds with uninformative priors constraining the various parameters as *P*(*a*) = *f_Unif_*(*a*|0,10000), *P*(*b*) = *f_Unif_*(*b*|0,10000), *P*(*δ*) = *f_Unif_* (*δ*|0.0001, 0.1), *P*(*κ*) = *f_Unif_* (*κ*|0,1000), *P*(*μ*) = *f_Unif_* (*μ*|0,100000) where

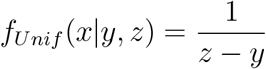

with support [*y,z*]. This first stage of inference allows estimation of *a* and *b* but not *δ*. A second stage of inference is then carried out on each gene, with information carried through from stage 1 for the initialisation by taking the 0.00005 and 0.99995 quantiles of *P*(*θ*|*L*) to obtain *a_min_*, *a_max_*, *b_min_*, *b_max_*, *μ_min_*, *μ_max_* under the assumption of normally distributed posteriors. Stage 2 relies on model 2, using equation 14 as the likelihood function (unless the hybdrid approach is used) to obtain the posterior *P*(*θ*|*Y, L*). Before initialisation, if the upper bound from equation 1, *M* ≥ 7000 and 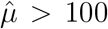 then the state space of the gene is determined to be too large for the analytical approach, and the the analytical/numerical hybrid approach is used instead. In this case, the *ϵ* value must first be calculated using a similar drawing scheme as described before for the initialisation to draw a sample of 1000 sets of *θ*. This time, draw

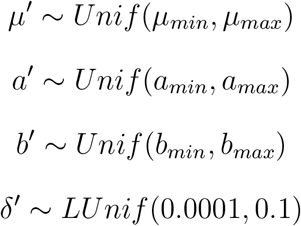

and set *b* = (*μ*’*a*’/*b*’)^1/2^, *a* = *ba*’/*b*’ and *δ* = *δ*’ for *θ* = (*a,b,δ*), repeating 1000 times to obtain the sample. Then simulate a dataset, *Y*’, for each sampled *θ*, with each comprising *N* = 795 simulated cells, and find the distances, *D*(*Y’, Y*). Then set *ϵ* = min(**D**) + 0.05(max(**D**) — min (**D**)), where **D** = (*D*_1_,…, *D*_1000_).

If the fully analytical approach is being used, the chain is initialised in the same manner as in stage 1 to draw *θ*_1_. If the hybrid approach is being used, draw using the same scheme that was used to obtain *ϵ*. The same uninformative priors as in stage 1 are used in stage 2 for both the analytical and hybrid approaches. If stage 2 is begun with the analytical approach, there may still be two instances that require switching to the hybrid approach. Firstly, if the chain converges to a region of parameter space which results in the state space being too large then the switch must be made. This is implemented by checking at each step if *M* ≥ 7000 or *M* ≥ 1000 ⋂ *a* < 1/1000 after the 500th step in the chain and switching if either is true. The other scenario occurs when the gene resides within unstable parameter space, resulting in negative probability values from equation 8. If a proposal is made to a point in unstable parameter space, it will be rejected on the basis of negative probability values. Therefore, we determine if the gene lies within unstable space by checking at each step if the rate of rejection due to an unstable proposal ≥ 0.05 for steps [max(1000, *j* – 5000), *j*] when *j* > 2000. In both cases, the chain is restarted using the hybrid approach, carried out in the exact same manner as before, except now simulating *N* = 7950 cells for both the *ϵ* calculation and during the actual chain progression, instead of the previously used *N* = 795. Inference with model 2 allows estimation of *a, b* and *δ*, recovering all parameters of interest from the 4sU scRNA-seq dataset.

In summary, we sampled two posterior distributions for each gene analysed, using model 1 and 2, to obtain *P*(*θ*|*L*) and *P*(*θ*|*Y, L*) (equations 5 and 16). Estimates for each parameter may then be derived as the mean of the posterior, *E*(*θ_i_*) = ∫ *θ_i_P*(*θ_i_*|*Y, L*)*dθ_i_*, with confidence in our estimates being quantified with the coefficient of variance, 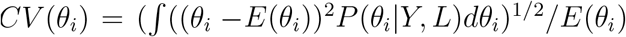. Inference on the data from Qiu et al. 2020 [38] was carried out for all 13264 genes which had at least one read and observed T>C conversion in both the 4sU and control datasets. Immediately 28 genes were removed whose model 2 Markov chain proposal acceptance rates were < 0.1 (SI), which represent those with very rough/discontinuous *D*(*Y*\*, *Y*) ~ *δ* due to having very low *μ* and therefore sparse data, leaving 13236 genes. This means that the hybrid approach was utilised in stage 2 of inference for these 28 genes but faced technical issues due to the sparseness of the data.

If a gene has a very low (close to background) or very high (close to the theoretical maximum) conversion rate in the 4sU dataset, it is not possible to reliably estimate δ. In such cases, neither the estimate or confidence quantification are reliable and therefore those genes had to be discarded (SI).

### Simulations for model comparison

The performance of inference using different likelihood functions was tested on simulated data. Gillespie’s exact algorithm (stochastic simulation algorithm) [65] was used to simulate data according to the input matrix shown in table 3 and the output matrix shown in table 4, with the stoichiometry matrix shown in table 5.

**Table 3:** Input matrix for new and surviving transcript count Gillespie algorithm simulations.

| | $RNA_0$ | $RNA_1$ | $G_{on}$ | $G_{off}$ |
| --- | --- | --- | --- | --- |
| $\beta_0$ | 0 | 0 | 1 | 0 |
| $\beta_1$ | 0 | 0 | 1 | 0 |
| $\delta_0$ | 1 | 0 | 0 | 0 |
| $\delta_1$ | 0 | 1 | 0 | 0 |
| $k_{on}$ | 0 | 0 | 0 | 1 |
| $k_{off}$ | 0 | 0 | 1 | 0 |

**Table 4:** Output matrix for new and surviving transcript count Gillespie algorithm simulations.

| | $RNA_0$ | $RNA_1$ | $G_{on}$ | $G_{off}$ |
| --- | --- | --- | --- | --- |
| $\beta_0$ | 1 | 0 | 1 | 0 |
| $\beta_1$ | 0 | 1 | 1 | 0 |
| $\delta_0$ | 0 | 0 | 0 | 0 |
| $\delta_1$ | 0 | 0 | 0 | 0 |
| $k_{on}$ | 0 | 0 | 1 | 0 |
| $k_{off}$ | 0 | 0 | 0 | 1 |

**Table 5:** Stoichiometry matrix for new and surviving transcript count Gillespie algorithm simulations.

| | $RNA_0$ | $RNA_1$ | $G_{on}$ | $G_{off}$ |
| --- | --- | --- | --- | --- |
| $\beta_0$ | 1 | 0 | 0 | 0 |
| $\beta_1$ | 0 | 1 | 0 | 0 |
| $\delta_0$ | -1 | 0 | 0 | 0 |
| $\delta_1$ | 0 | -1 | 0 | 0 |
| $k_{on}$ | 0 | 0 | 1 | -1 |
| $k_{off}$ | 0 | 0 | -1 | 1 |

This allows simulation of the pool of transcripts in the cell that was synthesised before (*RNA*_0_) and after (*RNA*_1_) the 4sU pulse started. The simulation is run with initial conditions *X*_0_ = (0, 0, 0,1) where *X* = (*RNA*_0_, *RNA*_1_, *G_on_*, *G_off_*) and rate constant values are set for bursty expression *θ* = (*β*_0_ = 50, *β*_1_ = 0, *δ*_0_ = 0.001, *δ*_1_ = 0.001, *k_on_* = 0.0005, *k_off_* = 1), running until *t*_0_ = 200000 to bring the system to steady state. The system state at the end of this run, *X*_*t*0_, is then used as the initial condition for a second run, where we now set *θ* = (*β*_0_ = 0, *β*_1_ = 50,*δ*_0_ = 0.001,*δ*_1_ = 0.001, *k_on_* = 0.0005, *k_off_* = 1) to simulate the newly synthesised transcripts produced during the 4sU pulse. A pulse duration of *t*_1_ = 1000 minutes was used here, giving the final state of the system *X*_*t*1_, and importantly giving the counts for *RNA*_0_ and *RNA*_1_ in the cell. This was repeated to simulate *N* = 10000 cells. In silico sequencing data was then generated based on these simulated transcript count values. Cell-specific capture efficiencies were drawn

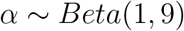

before drawing the cell-specific UMI counts, *l*, corresponding to the two pools of transcripts as

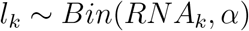

for *k* = 0 and *k* = 1, so that the total UMI count for the given cell is *l* = *l*_0_+*l*_1_. The cell-specific total number of reads corresponding to each UMI in the two pools is then drawn

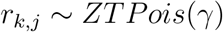

where *γ* = 5 represents sequencing depth and reads per UMI is a zero-truncated poisson random variable with

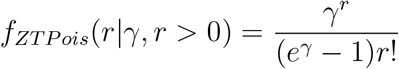

using the same logic of poisson assignment of reads to UMIs as in [66]. Then the cell-specific total number of reads of the given pool is

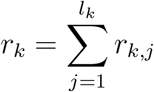

The number of uracils across the sequenced part of the transcript is then drawn for each read

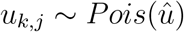

where *û* = 60 is the average number of uracils per read. The number of conversions in each read in the cell is then drawn for the two pools of transcripts as

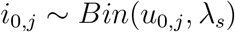

and

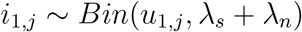

where we set λ_*s*_ = 0.01 and λ_*n*_ = 0.075. The overall conversion data across all reads in the cell is then *i* = (*i*_0_,*i*_1_), where *i*_0_ = (*i*_0;1_,…, *i*_0,*r*_0__) and *i*_1_ = (*i*_1;1_,…, *i*_1,*r*_1__), from which we obtain *y*, where *y_i_* is the number of reads with *i* conversions in the given cell. Now we have our simulated dataset which we can use to demonstrate our capacity to recover known parameter values. MCMC was carried out with fully analytical likelihood functions, although the chains were run only until 2000 steps were completed instead of the aforementioned 10000 steps. Posterior distributions were obtained from these Markov chains by taking the last 501 steps of the chain, using a thinning factor of 5, resulting in a sample size of 101.

## Supplementary information

### Cell-specific capture efficiencies

Our models require the capture efficiency, *α*, (proportion of transcripts from each cell with at least 1 corresponding read) of each cell to be known. This neccessitates the use of RNA spike-in probes, in which a known quantity of material is added to each cell and the proportion of molecules detected in the sequencing gives *α*. Spike-ins were not used in the Qiu datasets, but capture efficiencies may be inferred by using data from Klein et al 2015 [67], which has cell-matched (K562) scRNA-seq data (with GEO sample ID GSM1599501) which does contain ERCC spike-in probes. We construct a mathematical model to obtain the probability distribution of the capture efficiencies in the 4sU Qiu dataset, *α_q_* based on the Klein data, under the assumption that since both datasets were produced with K562 cells, the underlying probability distribution of the total transcript count in each cell, *m*, is the same for both datasets. According to [56] the true number of spike-in molecules loaded to each cell, *x*, may be modelled as a poisson random variable with rate based on the expected number of molecules loaded per cell, *λ*, (12467.64 in the case of the Klein dataset)

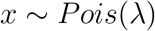

The capture efficiency of each of the 953 cells in the Klein dataset, *α_k_*, is then

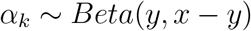

where *y* represents the total number of spike-in molecules detected in the given cell. The total number of transcripts present in each cell in Klein is modelled as

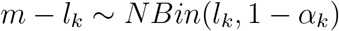

where *l_k_* represents the total UMI counts across all genes in Klein for the given cell, using the negative binomial parametrisation defined for equation 1. The capture efficiency of each cell in Qiu, *α_q_*, may then be obtained using the total number of UMIs in the given cell across all genes, *l_q_*,

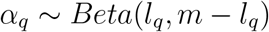

The probability density function for each cell in Qiu is then solved by numerically integrating the above distributions through random sampling to obtain

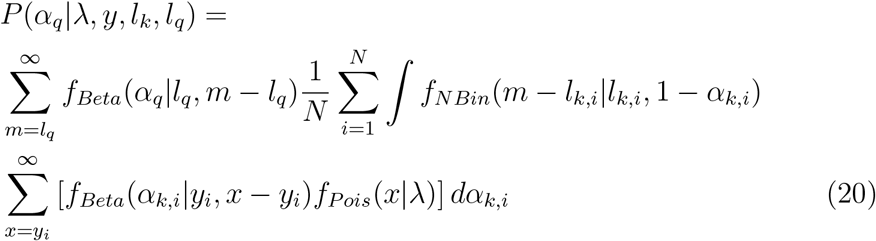

where *N* is the number of cells in Klein, *i* refers to the *i*th cell of Klein and

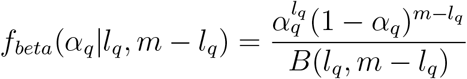

with

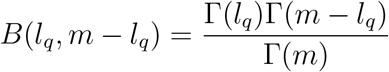

Estimates may be derived from *P*(*α_q_*|*λ,y,l_k_, l_q_*) with *E*[*α_q_*] and confidence may be quantified with 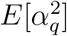. Figure 12 indicates high confidence in our estimates through the low CV, while the estimated capture efficiencies for Qiu are lower than for Klein, at around 0.02 on average. A quick, simple method for calculating *α_q_* estimates without quantifying confidence is as follows

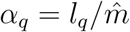

where

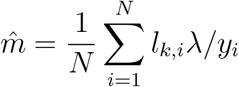

**Figure 12:**
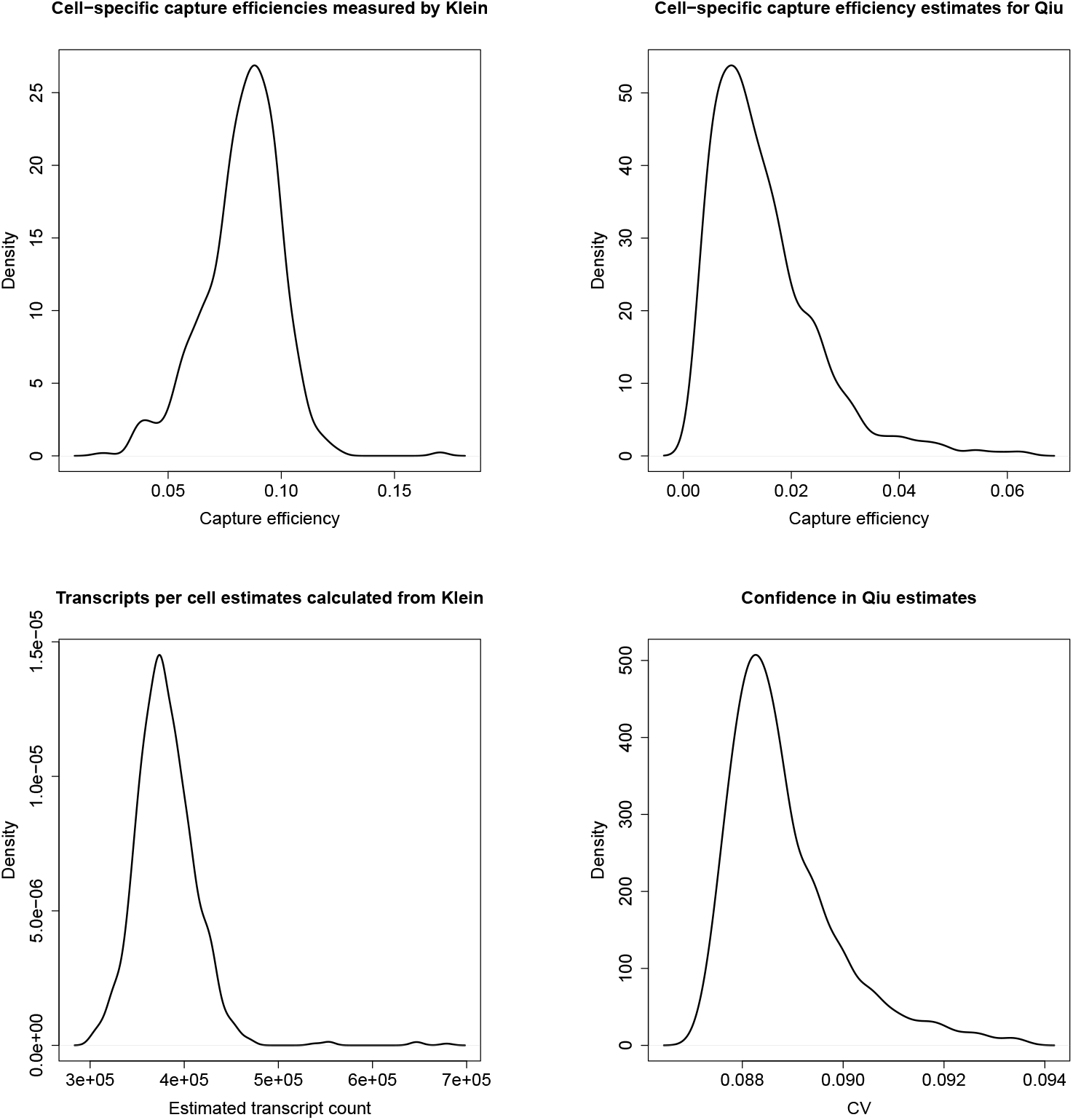
Density plots of the capture efficiencies measured for the Klein dataset, the total transcript content per cell estimated from Klein, the capture efficiencies estimated for the Qiu dataset and the confidence in those estimates as represented by the CVs.

### Conversion rates

Model 2 also requires the gene-specific background and gene-invariant 4sU-mediated T>C conversion rates to be known, λ_*s*_ and λ_*n*_, respectively. As previously mentioned, λ_*s*_ is defined as the proportion of genomic Ts in all reads and all cells that appeared as Cs in the control dataset for the given gene. Therefore, the conversion rate observed in the 4sU dataset corresponds to λ_*s*_ + λ_*n*_. The conversion rates of all genes for which we have high confidence in the rate estimate in both datasets are shown in figure 13, which was 4291 genes. Confidence is obtained by modelling the T>C rate, λ, as

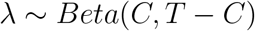

classing those with a resulting CV < 10^−0.5^ as having high confidence. We expect the rate in the 4sU dataset to be at least as large as in the control, hence genes tend to appear on the diagonal or above it. Genes with higher turnover are expected to appear further above the diagonal while those with low turnover are expected to appear closer to it. The curve along the top of the plot represents λ_*s*_ (x-axis) added to our estimate of λ_*n*_. λ_*n*_ is estimated by first assuming that all reads correspond to new transcripts (synthesised during the pulse) and then calculating

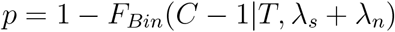

for each gene so that **p** = (*p*_1_, …,*p*_4291_) where

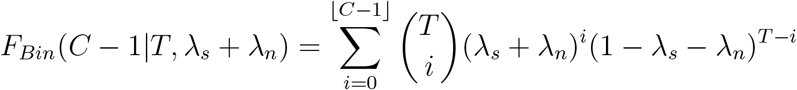

**Figure 13:**
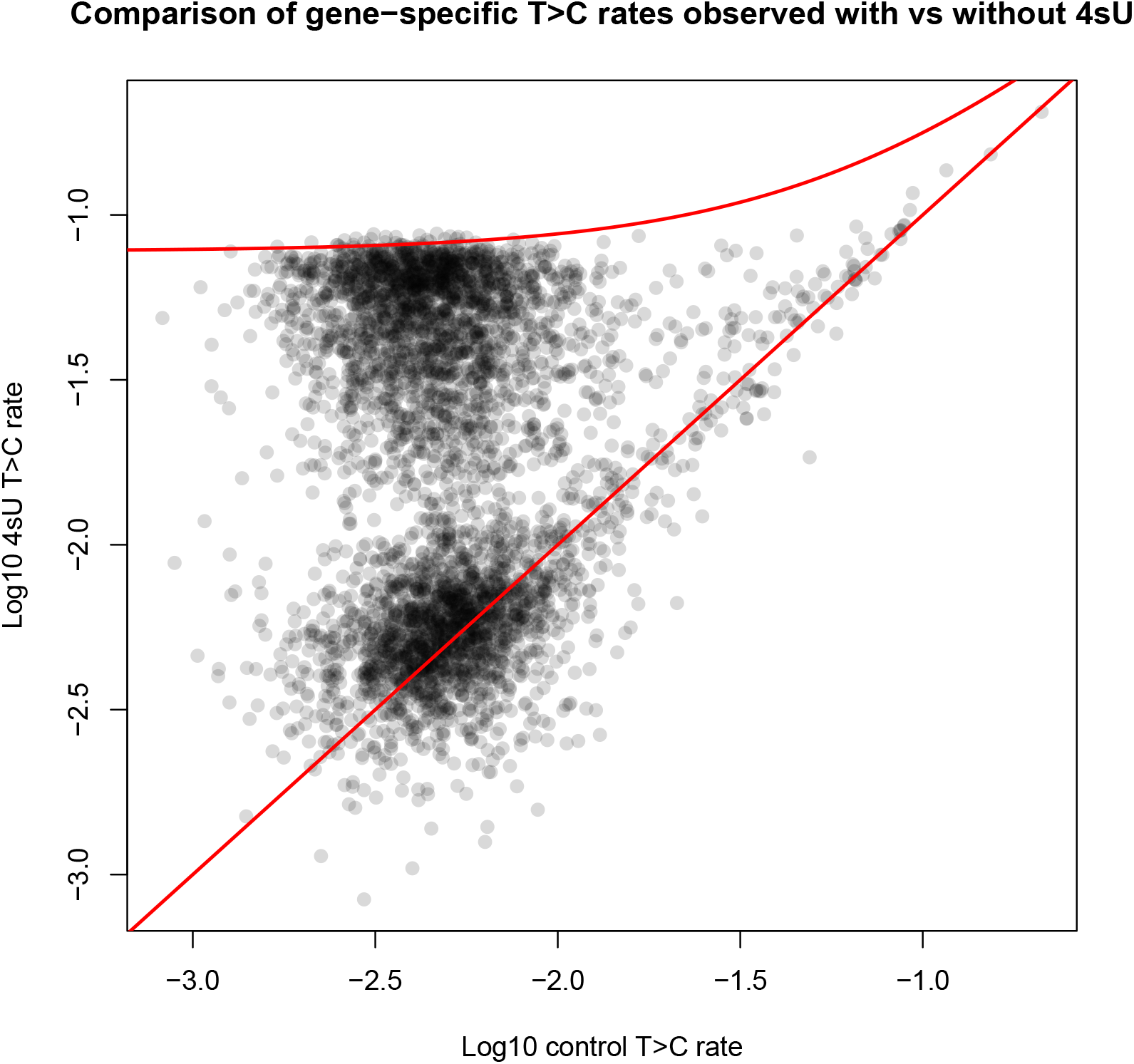
Gene-specific T>C conversion rates in the control vs 4sU datasets for 4291 genes for which we have high confidence in the observed T>C rate in both datasets. The lower red line represents λ_*s*_, while the upper one represents λ_*s*_ + λ_*n*_.

The estimate for λ_*n*_ is then the minimum value for which min(**p**) ≥ 1/4291. With this approach, λ_*n*_ ≈ 0.0775.

### Additional information on decay rate estimates

Uninformative uniform priors covering the entirety of plausible parameter space were used during the inference. However, if a gene has a very low (close to background) or very high (close to the theoretical maximum) conversion rate in the 4sU dataset, then it is not possible to reliably estimate *δ*. In such cases, the Markov chain will be pushed to ever lower or higher values along the *δ* axis and as a result will be “pinned” close to the lower or upper bound of *P*(*δ*), resulting in an artificially reduced CV for our estimate of *δ*. There is an association between the estimate and CV, in which genes with a transcript lifetime similar to the 4sU pulse duration used in the experiment (240 minutes), with log_10_(*δ*) ≈ –2.38, result in estimates with the highest confidence (lowest CV). CV increases for those genes with higher or lower estimates bidirectionally, until the aforementioned technical issue occurs for genes with unmeasurable *δ* where the CV decreases again as the chains drift towards the prior boundaries. The CV must be used to select for those genes with high confidence parameter estimates. Therefore, those genes with artificially reduced CV were discarded by accepting only those genes with log_10_(*E*(*δ*)) ∈ [–3.3, ‒1.7], as shown by the red lines in figure 14.

**Figure 14:**
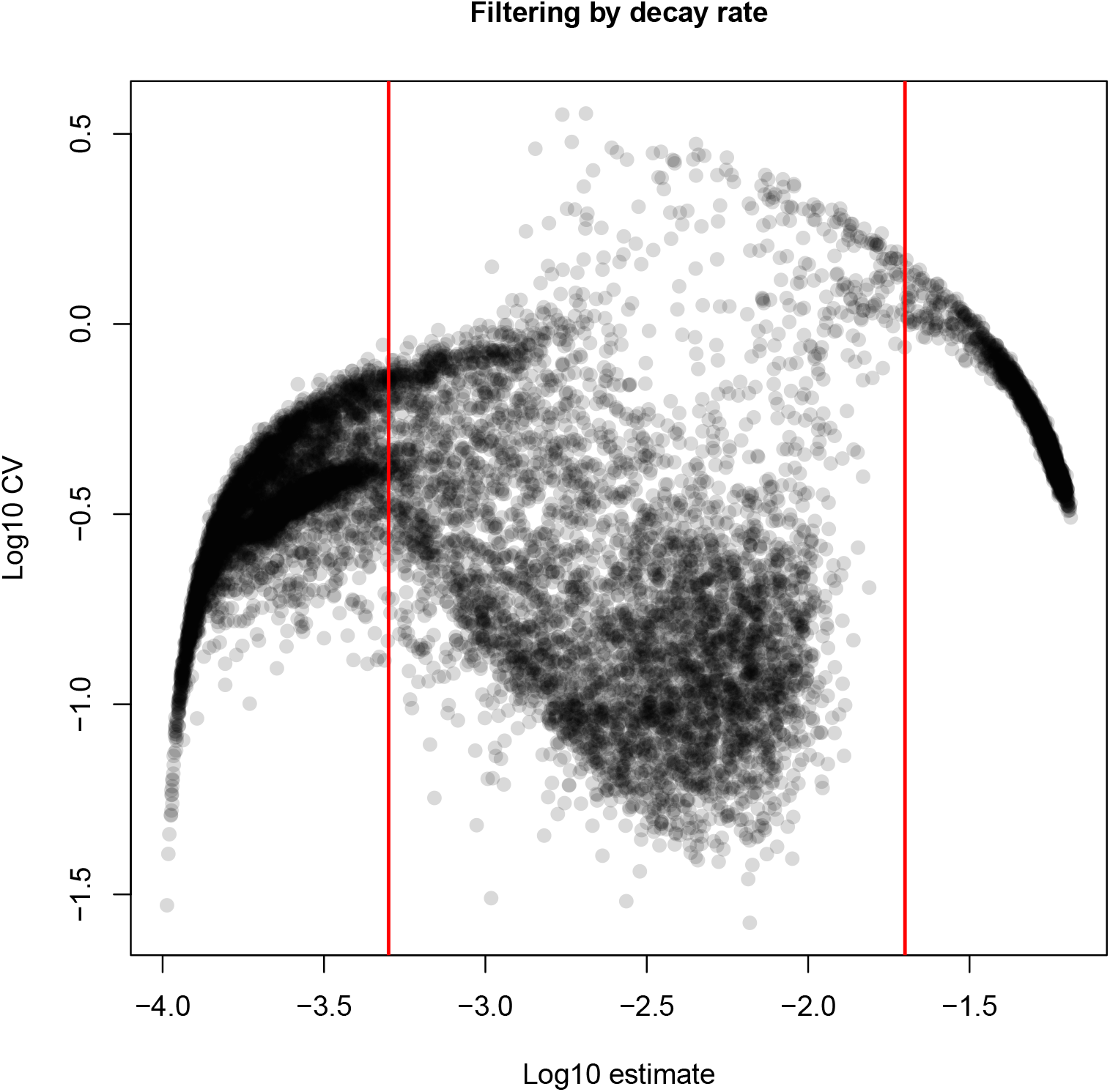
Decay rate estimates vs CVs obtained from stage 2 posteriors for 13236 genes, with imposed thresholds shown in red.

Additional confidence in our results is provided showing the strong correlation between the decay rate estimates we obtained from Qiu for our high confidence genes and those previously calculated in [36] for the same cell type (figure 15).

**Figure 15:**
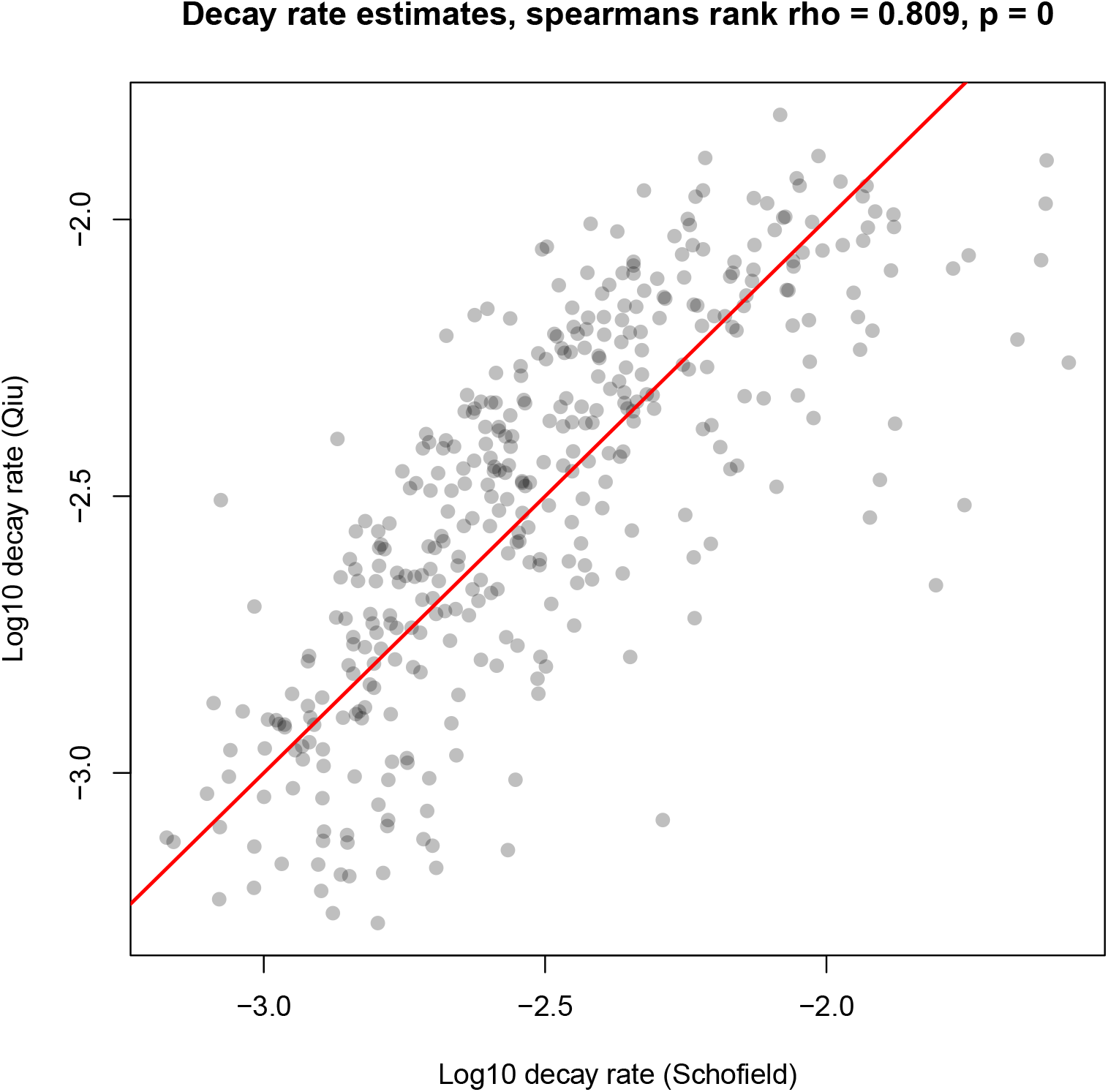
Decay rate (*δ*) estimates we obtained from Qiu for our high confidence gene set vs those calculated by Scofield for the same genes, with the Spearman’s rank correlation statistics shown and the diagonal displayed (red line).

### Inference on simulated data

In order to assess the performance of the inference algorithm, a dataset was simulated for each of the 13264 genes from the real Qiu dataset whose bursting dynamics were inferred. The *θ* estimates obtained in stage 2 of inference on Qiu were used as the “ground truth” parameter values to simulate new datasets. The simulation method is the same as that described for the hybrid inference approach (Methods) for simulation of surviving, *s*, and new, *n*, transcripts of each cell, but differs in the simulation of the T>C data since the single cell aspect is not neglected, and also includes simulation of the UMI count data. The number of UMIs corresponding to surviving and new transcripts in each cell is drawn as

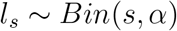

and

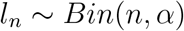

where *α* is sampled without replacement from the set of estimated capture efficiencies. We use *N* = 795 cells in the simulated data, matching the number of cells in the real dataset. Then we have the total UMI count for each cell *l* = *l_s_* + *l_n_* and *L*, where *L_c_* is the UMI count of cell *c*. For each UMI, we draw the number of corresponding reads, *r*, as

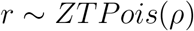

where *ρ* is the maximum likelihood estimate given the observed ratio of reads to UMIs, *R*, across all cells for the given gene in the real data. *ρ* is obtained by minimising

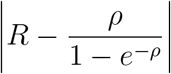

From this we have the total number of reads corresponding to surviving, *r_s_* and new, *r_n_* transcripts for each cell. The number of uracils, *u*, in each read is then drawn from the gene-specific observed probability mass function, *P*(*u*), from equation 11. Finally, the number of uracils in each read which undergo a T>C conversion, *i*, is then drawn for surviving reads as

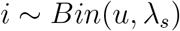

and for new reads as

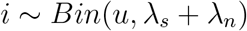

Now we have *y*, where *y_i_* represents the number of reads with i total conversions in the given cell and YC is the the vector *y* for cell *c*. We may now carry out inference with *L* and *Y* as previously described (Methods). The same filtering process that was applied to the real data was used here. The correlations between the parameter estimates derived from the stage 2 posteriors for the 603 filtered simulated genes are shown in figure 16. The strong, tight correlations along the diagonal demonstrate the successful recovery of ground truth parameter values for all parameters. The error increases for genes with very low *b* or *δ*, which reflects the increased CV for such estimates shown in figure 5. High error for *a* estimates primarily corresponds to those genes with very low, error-prone *b* values.

**Figure 16:**
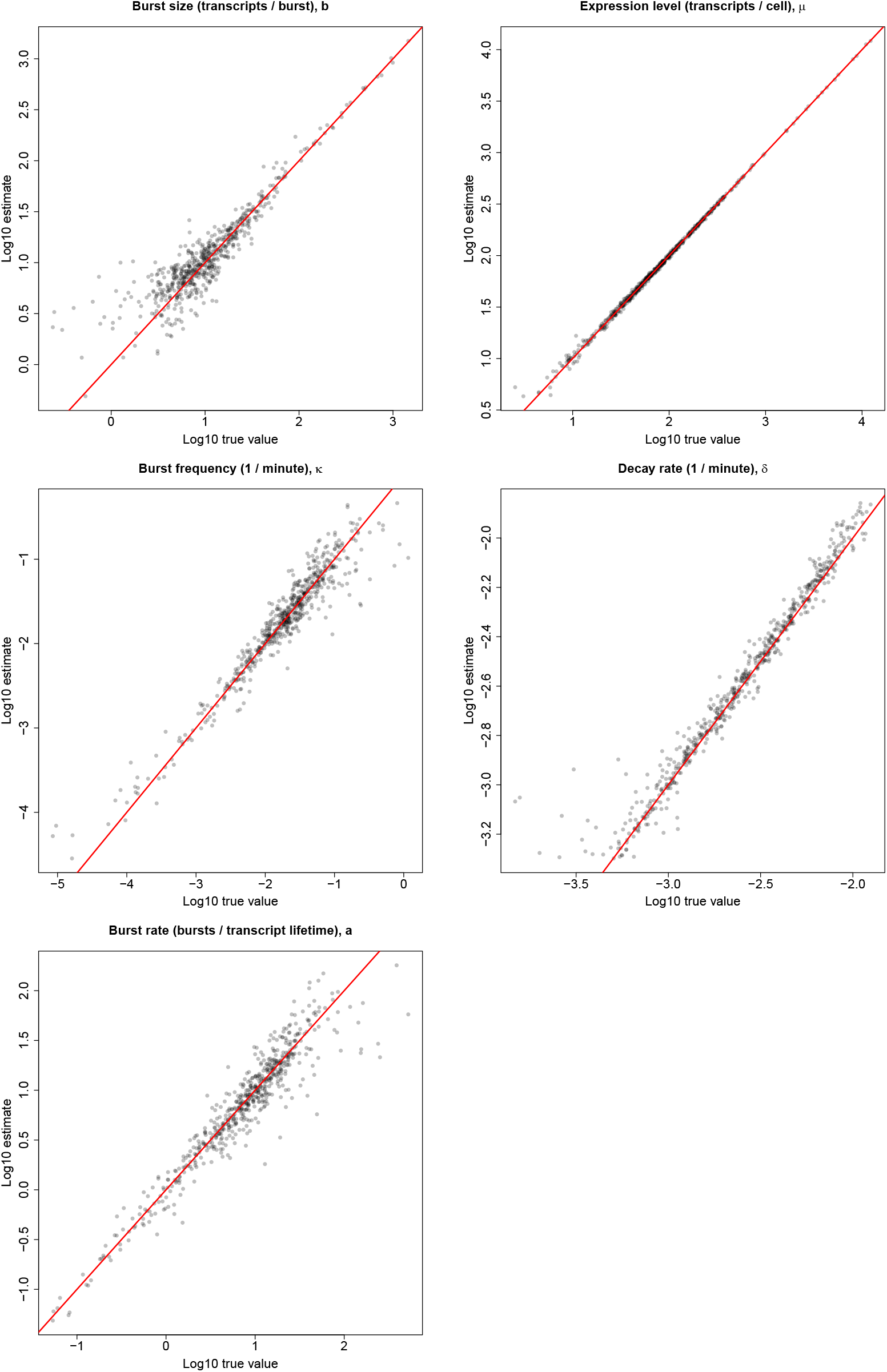
Correlations between true parameter values and those estimated from stage 2 posteriors for 603 simulated genes, with red lines representing diagonals.

### Acceptance rates

Density plots of the MCMC acceptance rates (after burn-in removal) for inference with both models 1 and 2 for all 13264 genes analysed in the Qiu dataset and the corresponding simulated genes are shown (figure 17). This diagnostic indicates that overall the desired mixing behaviour is achieved in all cases, with the ideal acceptance rate being roughly 0.234 [63].

**Figure 17:**
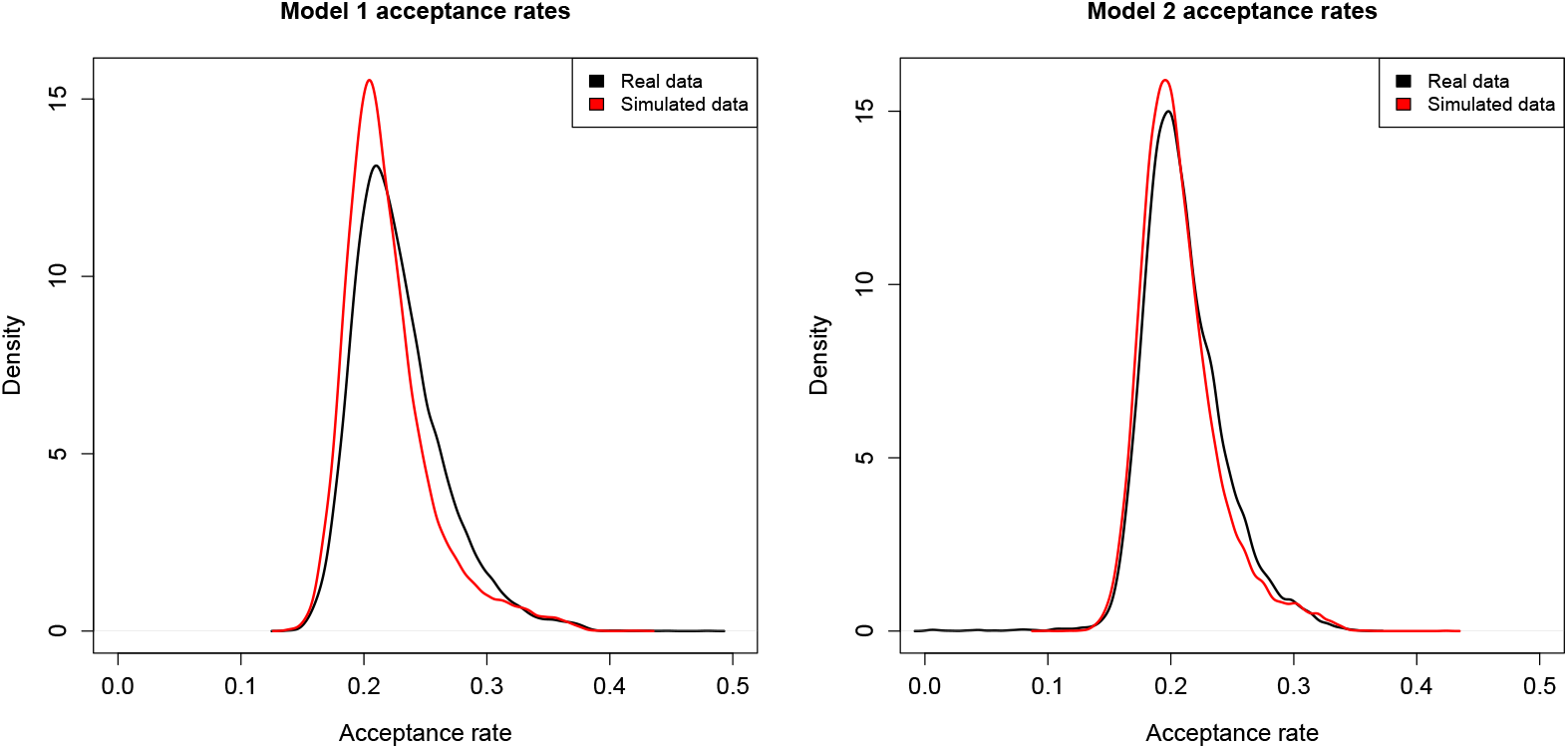
Markov chain acceptance rates (excluding burn-in) for both model 1 and 2 inference on the Qiu dataset and the corresponding simulated dataset.

### Further metagene and correlation analyses

#### Promoter-localised histone modifications

Metagene analyses of the other promoter-localised HMs that were represented by H3K4me2 (Results), showing the profiles for H3K4me3 (figure 18), H3K9ac (figure 19) and H3K27ac (figure 20).

**Figure 18:**
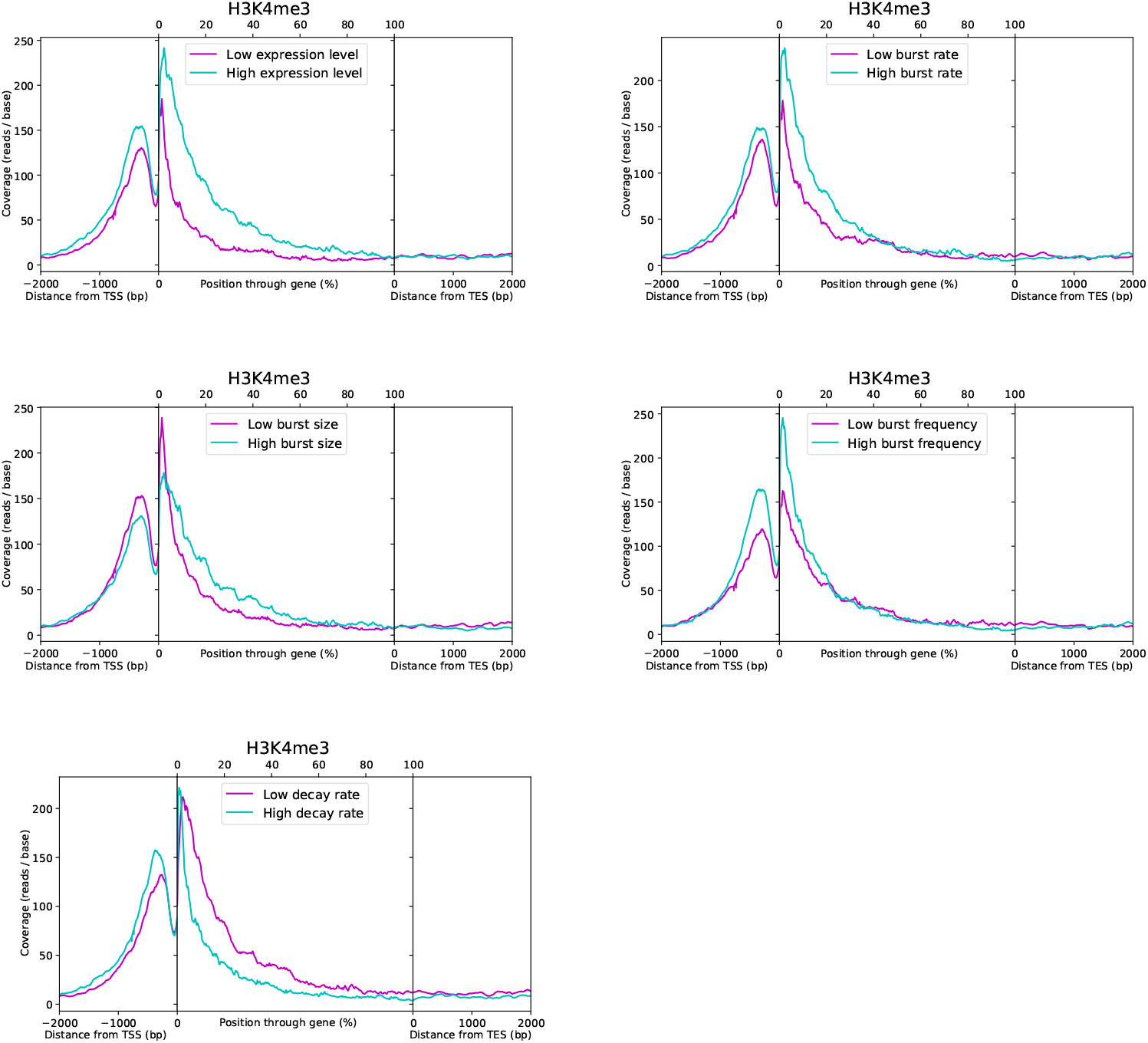
Metagene plots of H3K4me3 coverage, comparing profiles for the top and bottom 50% of genes when split according to their estimates for each parameter, denoted by high and low, as indicated.

**Figure 19:**
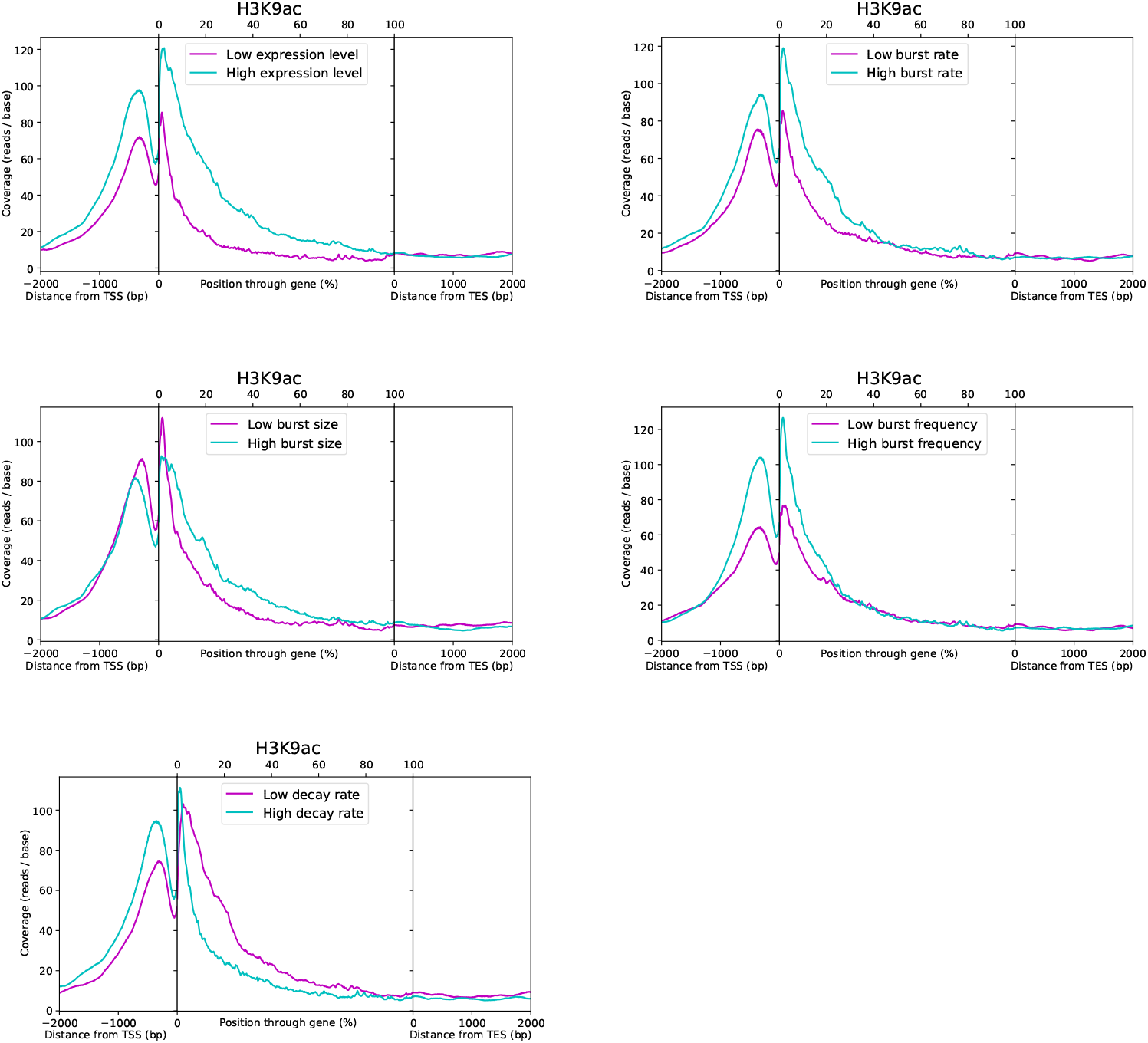
Metagene plots of H3K9ac coverage, comparing profiles for the top and bottom 50% of genes when split according to their estimates for each parameter, denoted by high and low, as indicated.

**Figure 20:**
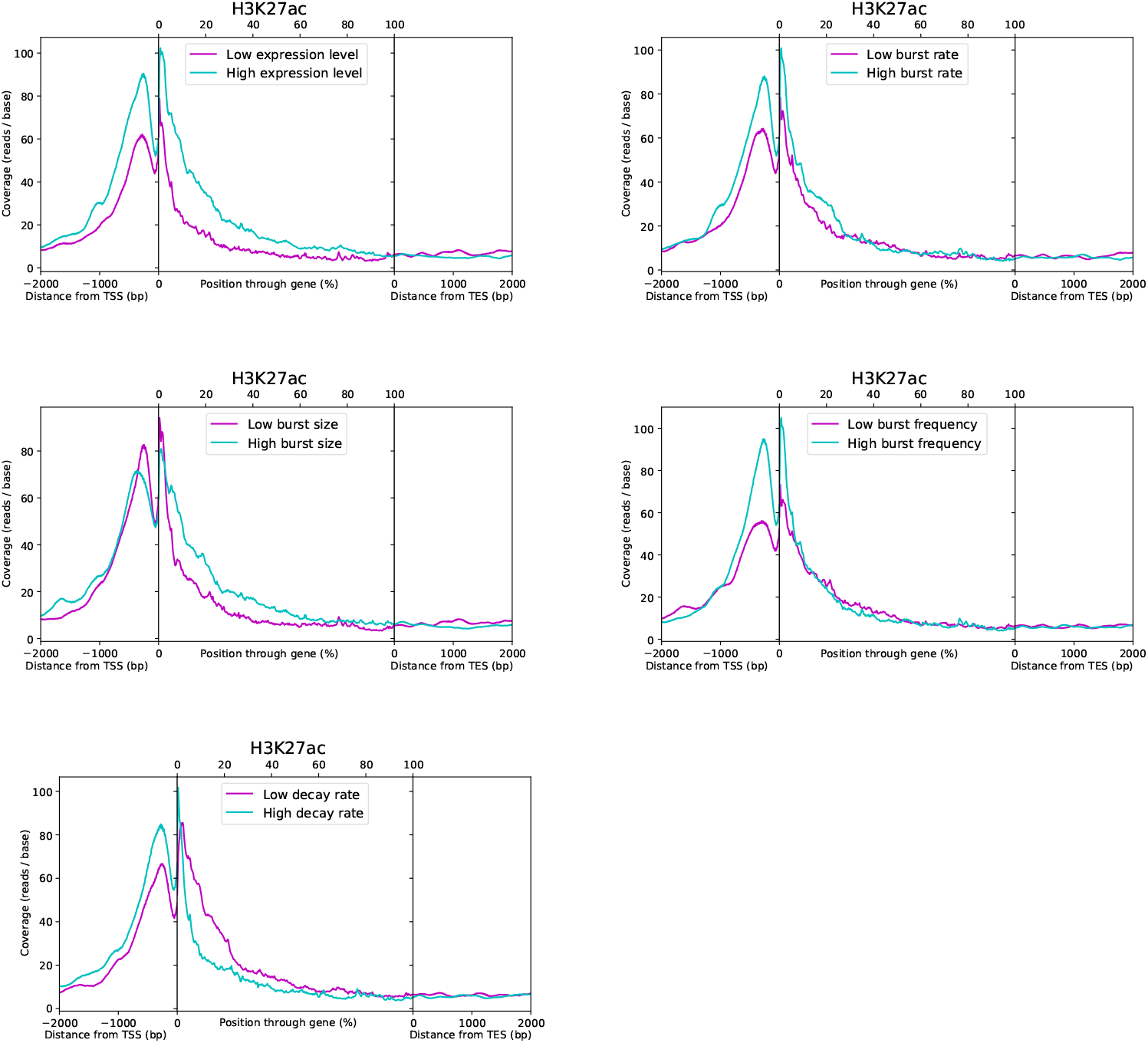
Metagene plots of H3K27ac coverage, comparing profiles for the top and bottom 50% of genes when split according to their estimates for each parameter, denoted by high and low, as indicated.

#### Gene body-localised histone modifications

A similar analysis of the GB-localised HMs was also carried out, where we use H3K36me3 as a representative example, although their metagene profiles and bursting associations are somewhat more diverse than with the four promoter-localised HMs. H3K4me1 was categeorised as being primarily promoter associated in [20] but we find its connection to bursting instead to be contingent upon its presence throughout the gene body (GB), and have therefore reclassified it for this context. The profiles of H3K36me3 halved by the different bursting parameter estimates as before (figure 21) indicate that presence throughout the GB and around the TES seems to be associated with increased *μ, a* and *κ* in a uniform manner. No association with *b* or *δ* is apparent, suggesting that this HM is associated with increased μ purely through increased *κ*. In this case, we are able to support the previously reported correlation with *a* [20], and confirm that the inability of scRNA-seq data to distinguish *a* and *κ* did not skew the final conclusions by quantifying the strength (figure 22) and statistical significance (figure 23) of H3K36me3 and the other GB-localised HMs which it represents (H3K79me2 and H4K20me1). It should be noted, however, that based on the metagene analysis, while both H3K79me2 and H4K20me1 appear to be primarily associated with increased *μ, a* and *κ*, along with H3K4me1, they look to have a positive and negative association with *b* and *δ*, respectively, when bound throughout the 20%:100% region, although this is statistically significant only for H3K4me1 (figure 23), which also has no association with *κ* and no significant correlation with a. Therefore, for the GB-localised HMs, H3K36me3, H3K79me2 and H4K20me1 can be regarded as similar in their associations with bursting dynamics, while H3K4me1 is an outlier. The regions of association for the GB-localised HMs vary to a degree, as dictated by the metagene analysis, with the values used for the correlation analysis shown in figures 22 and 23 being averaged across 0%:2000 (TSS to 2000bp downstream of the TES) for H3K36me3 and H4K20me1, −2000:100% for H3K79me2 and 20%:100% for H3K4me1. The metagene analyses of H3K4me1 (figure 24), H3K79me2 (figure 25) and H4K20me1 (figure 26) are shown.

**Figure 21:**
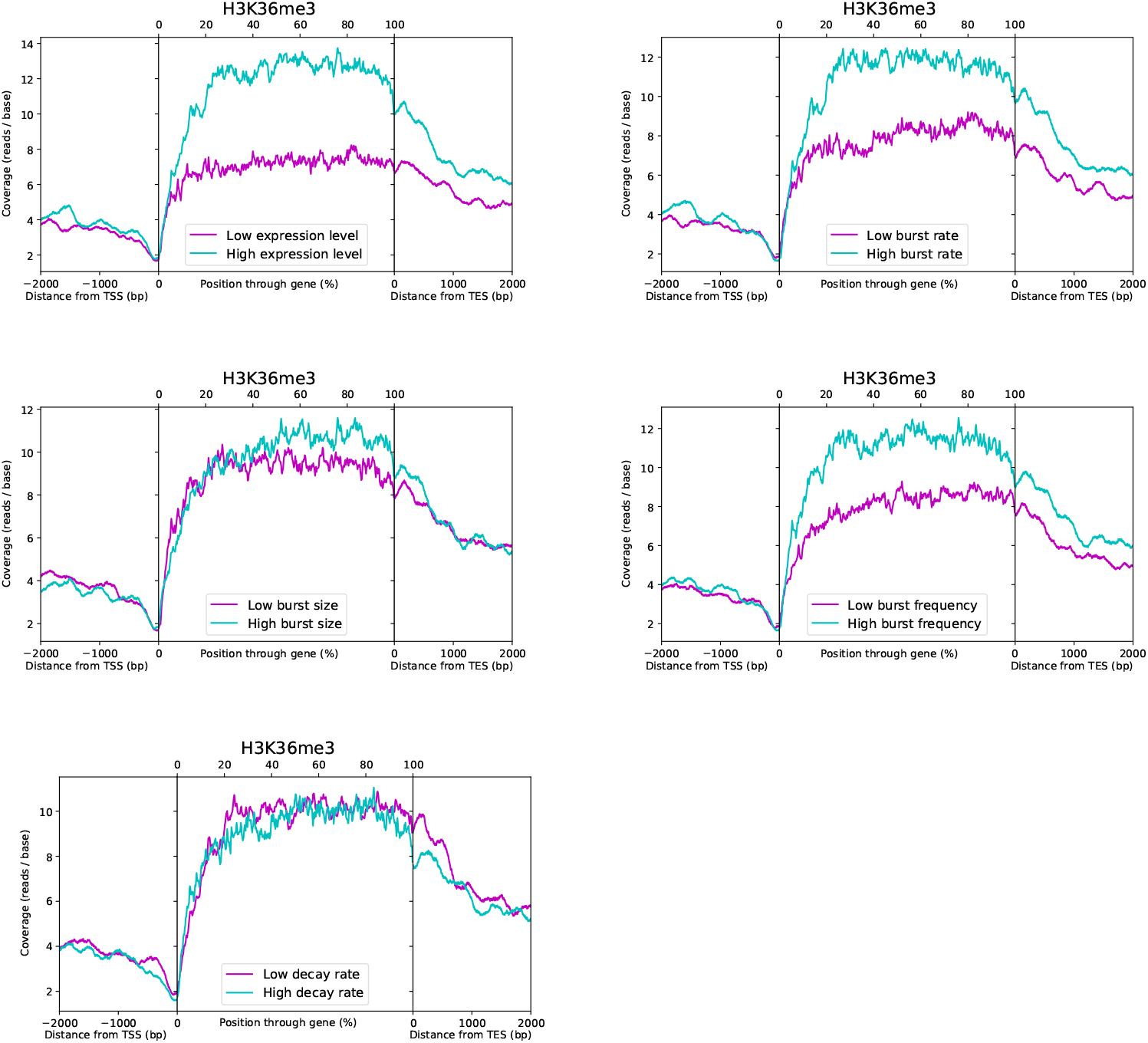
Metagene plots of H3K36me3 coverage, comparing profiles for the top and bottom 50% of genes when split according to their estimates for each parameter, denoted by high and low, as indicated.

**Figure 22:**
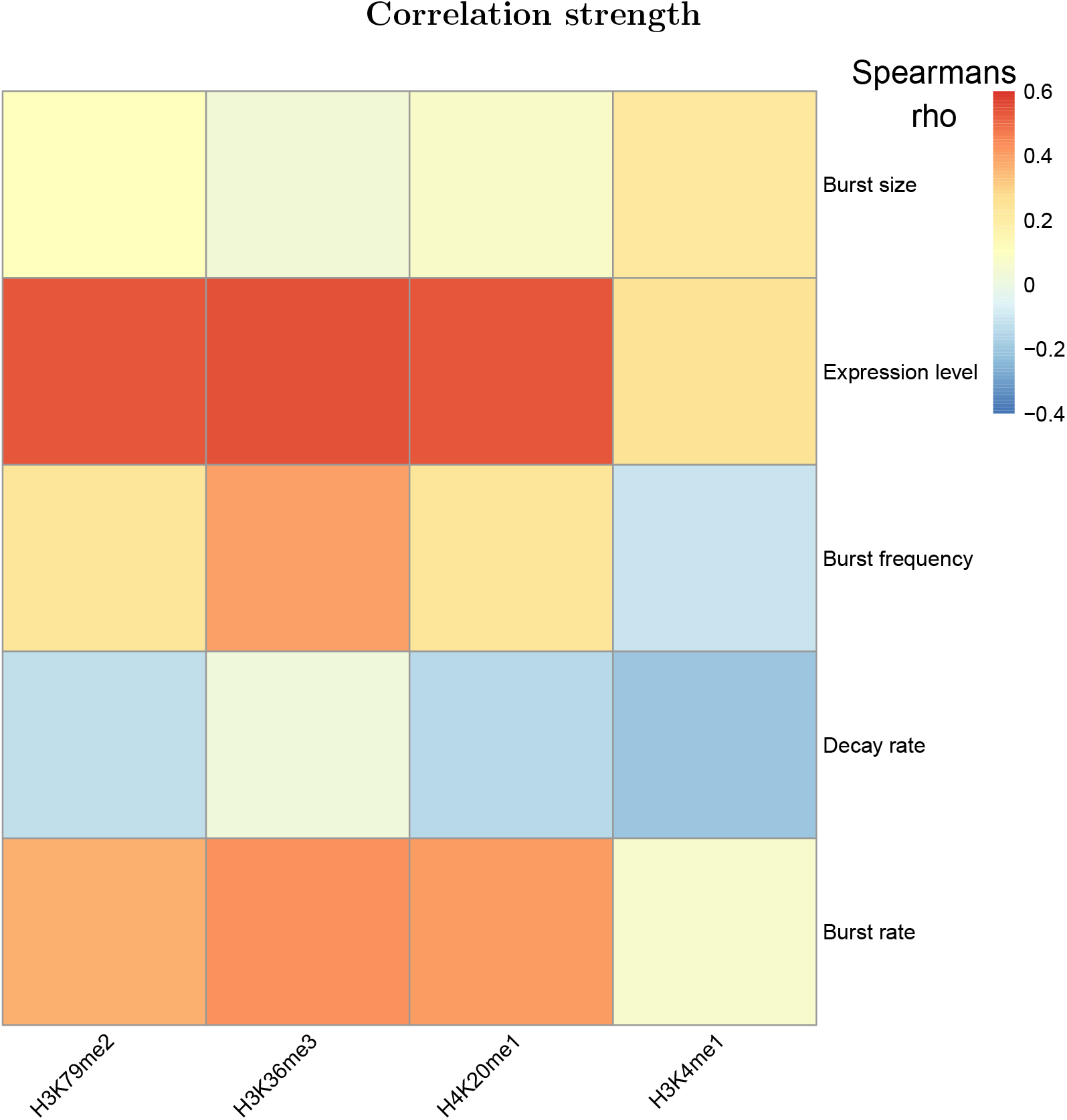
Heatmap showing the Spearman’s rank rho as the heat intensity value for the correlations between bursting parameter estimates and the mean GB-localised HM coverage values across 0%:2000 for H3K36me3 and H4K20me1, −2000:100% for H3K79me2 and 20%:100% for H3K4me1. More intense red or blue colouration indicates a stronger positive or negative correlation, respectively, while neutral indicates no/weak correlation.

**Figure 23:**
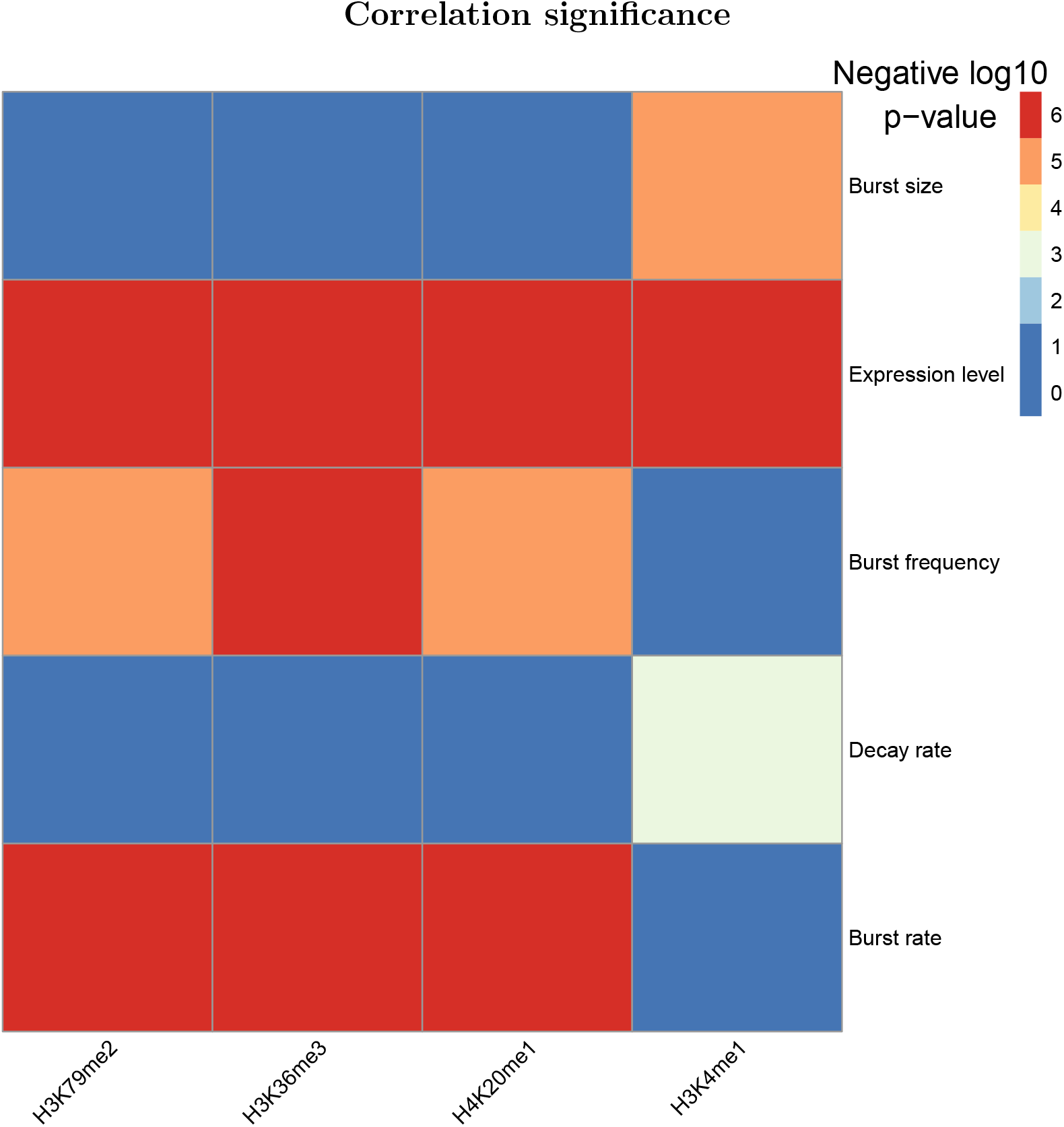
Heatmap showing the Spearman’s rank p-value (adjusted for multiple hypothesis testing) as the heat intensity value for the correlations between bursting parameter estimates and the mean GB-localised HM coverage values across 0%:2000 for H3K36me3 and H4K20me1, −2000:100% for H3K79me2 and 20%:100% for H3K4me1. The heat values are discretised, corresponding to negative log10 p-value thresholds. For example, the most intense blue indicates that, for the given correlation, 10^-2^ < *p*, meaning no statistical significance, the neutral colour indicates that 10^-4^ < *p* ≤ 10^-3^, while the most intense red indicates that *p* ≤ 10^-6^.

**Figure 24:**
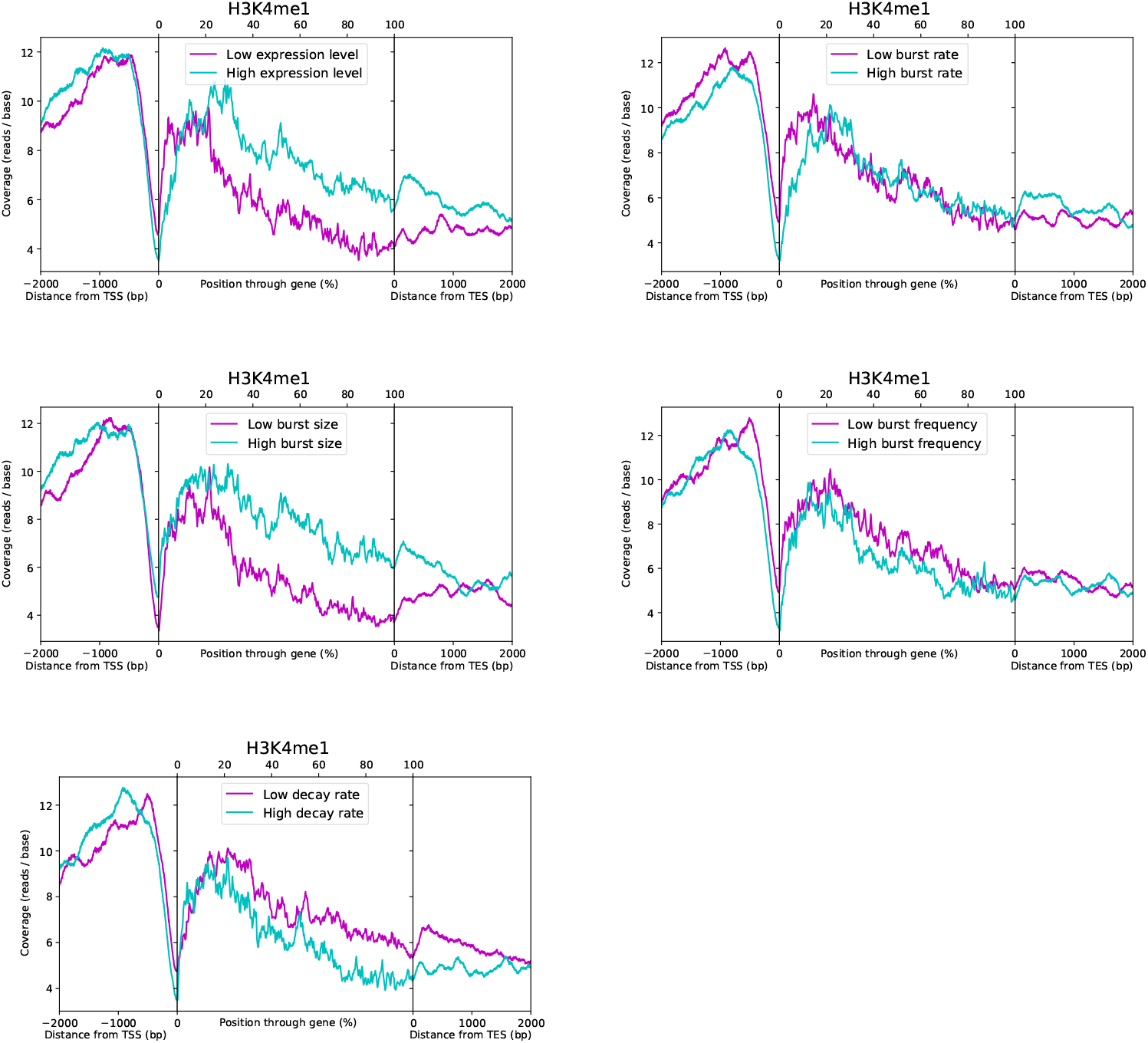
Metagene plots of H3K4me1 coverage, comparing profiles for the top and bottom 50% of genes when split according to their estimates for each parameter, denoted by high and low, as indicated.

**Figure 25:**
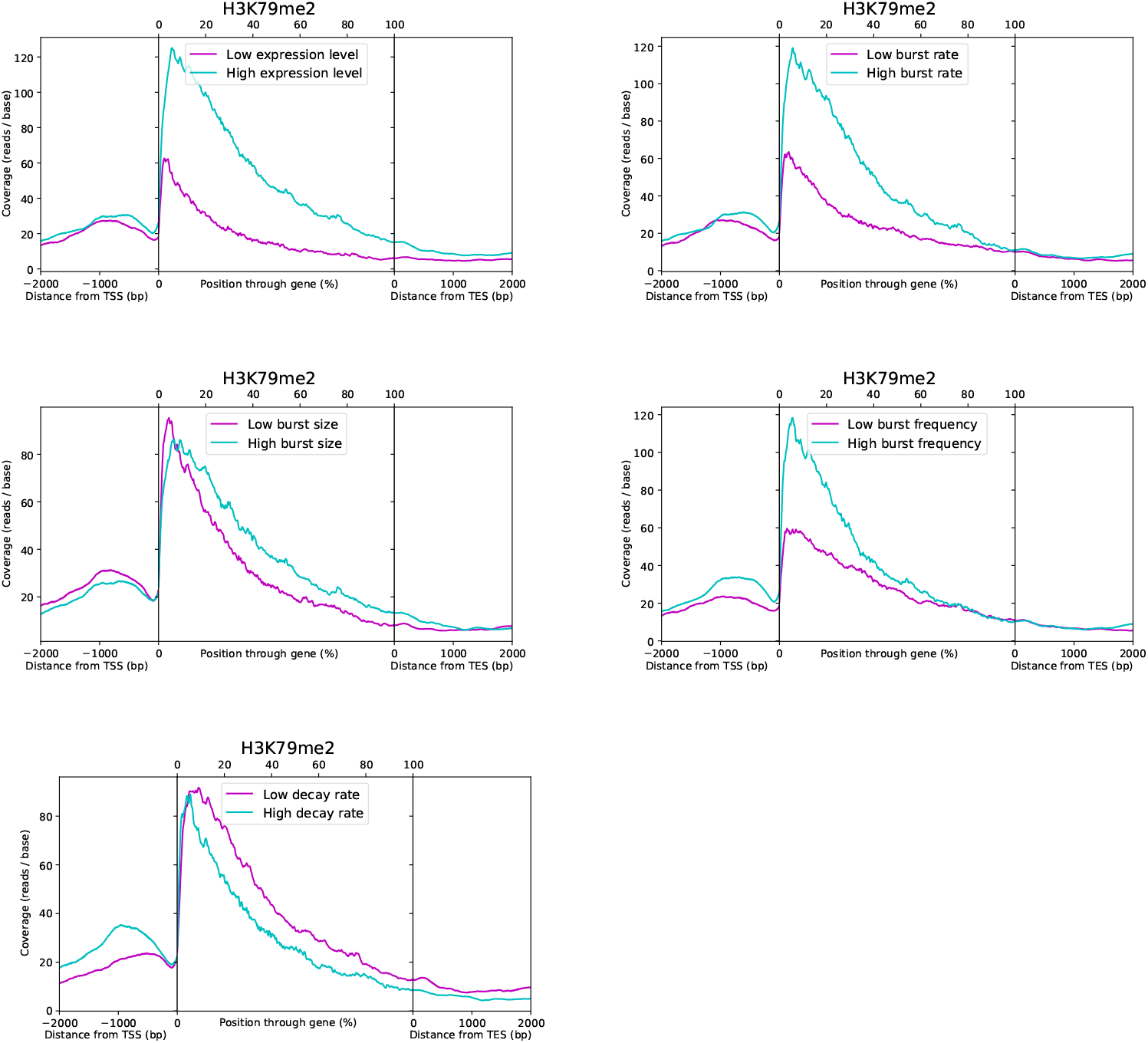
Metagene plots of H3K79me2 coverage, comparing profiles for the top and bottom 50% of genes when split according to their estimates for each parameter, denoted by high and low, as indicated.

**Figure 26:**
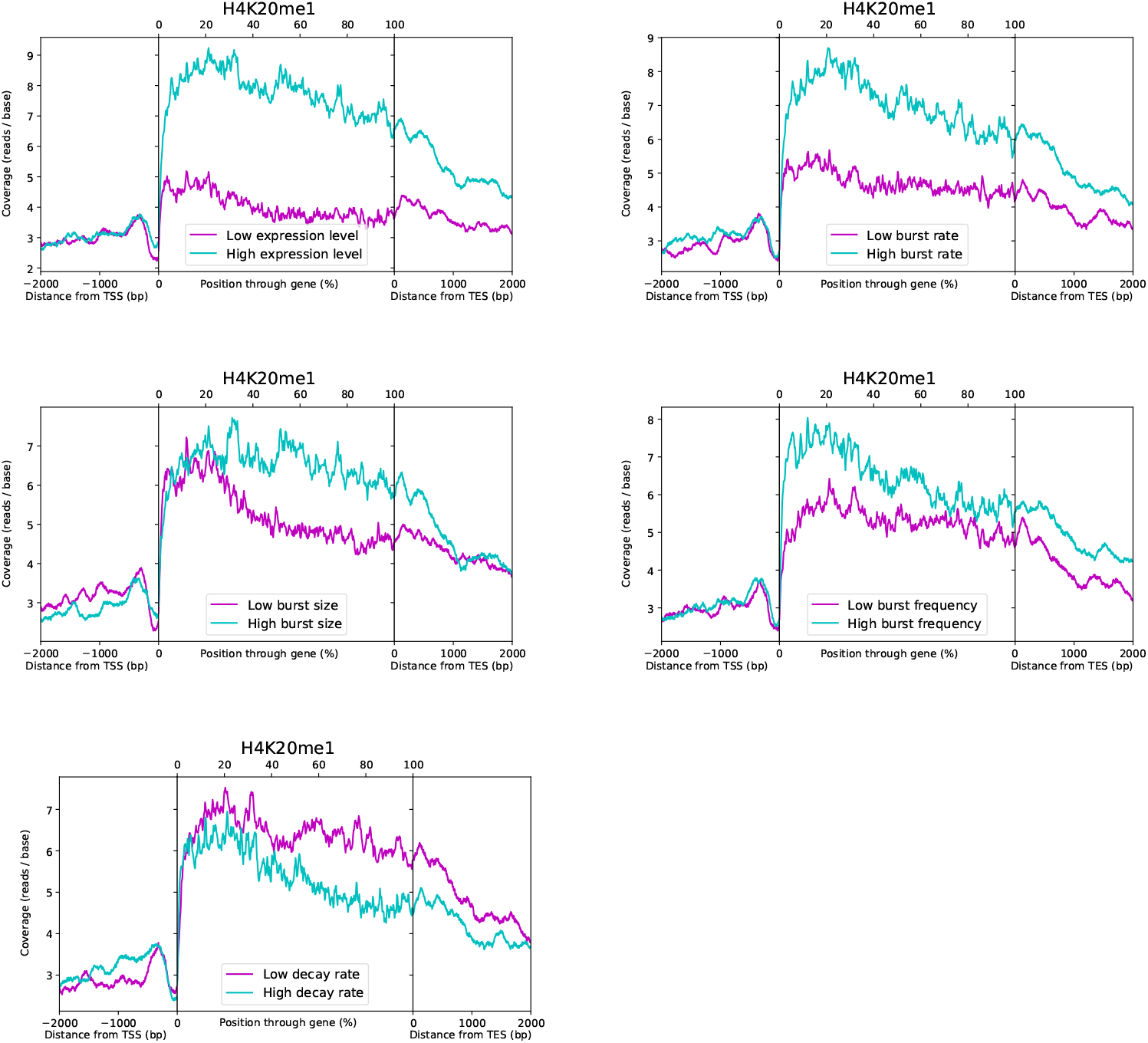
Metagene plots of H4K20me1 coverage, comparing profiles for the top and bottom 50% of genes when split according to their estimates for each parameter, denoted by high and low, as indicated.

### Additional Files

#### Additional file 1

Video gif showing the differential transition from surviving to new transcript pool for high and low noise genes through the cell-specific T>C rate distributions for data simulated with different pulse durations.

## Declarations

## Acknowledgements

We thank Louise Dyson for useful discussions regarding the mathematical theory relevant to the study and Francesca Mantellino for valuable input on the data processing strategy.

## Availability of data and materials

Parameter estimates and confidences obtained from the published 4sU scRNA-seq data will be deposited to an online repository like GEO and the code for the inference algorithm will be made available in a GitHub repository.

## Authors’ contributions

DME, PD and DH conceived the study while DME and DH designed the work. DME and PD acquired the data which DME analysed and DME, PD and DH interpreted. DME and PD created the software. DME wrote the manuscript with input from PD and DH. All authors approve the submission of the manuscript.

## Ethics approval and consent to participate

Not applicable

## Consent for publication

Not applicable

