## Supplementary figures and images for "Synergising single-cell resolution and 4sU labelling boosts inference of transcriptional bursting"

### Additional File 1

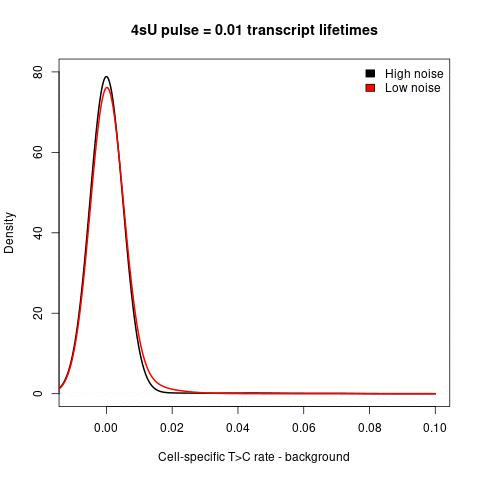
